# Orai3 orchestrates gemcitabine resistance in pancreatic cancer via NFATc1-SLIT3 axis

**DOI:** 10.1101/2025.08.22.671673

**Authors:** Samriddhi Arora, Abhishek Tanwar, Gyan Ranjan, Rajender K Motiani

## Abstract

Pancreatic Cancer (PC) is one of the most aggressive cancers and is associated with poor prognosis. Gemcitabine is the first-line chemotherapy for PC. While gemcitabine-based regimes offer survival benefits, the acquired gemcitabine resistance leads to reoccurrence, metastasis and the long-term survival rate remains dismal. Although there is substantial clinical evidence for gemcitabine resistance, the cellular and molecular mechanisms driving gemcitabine resistance remain largely unappreciated. Here, we reveal that Orai3, a Ca^2+^ selective channel, is a crucial driver of gemcitabine resistance. We demonstrate that Orai3 is upregulated in gemcitabine-resistant PC cells. Orai3 silencing in these cells decreases proliferation, induces cell cycle arrest, enhances apoptosis, and moderates stemness characteristics. Notably, studies in zebrafish model corroborate the significance of Orai3 in gemcitabine resistance *in vivo*. Mechanistically, our unbiased RNA-seq analysis coupled with robust functional studies show that SLIT3 works downstream of Orai3 to drive gemcitabine resistance. Finally, we report that NFATc1 transcription factor bridges Orai3 to SLIT3 transcription. Taken together, this study identifies Orai3 as a key orchestrator of gemcitabine resistance and uncovers a unique Orai3-NFATc1-SLIT3 signaling module that drives chemoresistance. Hence, this work reveals Orai3 as a promising target for synergistic therapeutic approach to combat chemoresistance.

**Graphical abstract:** 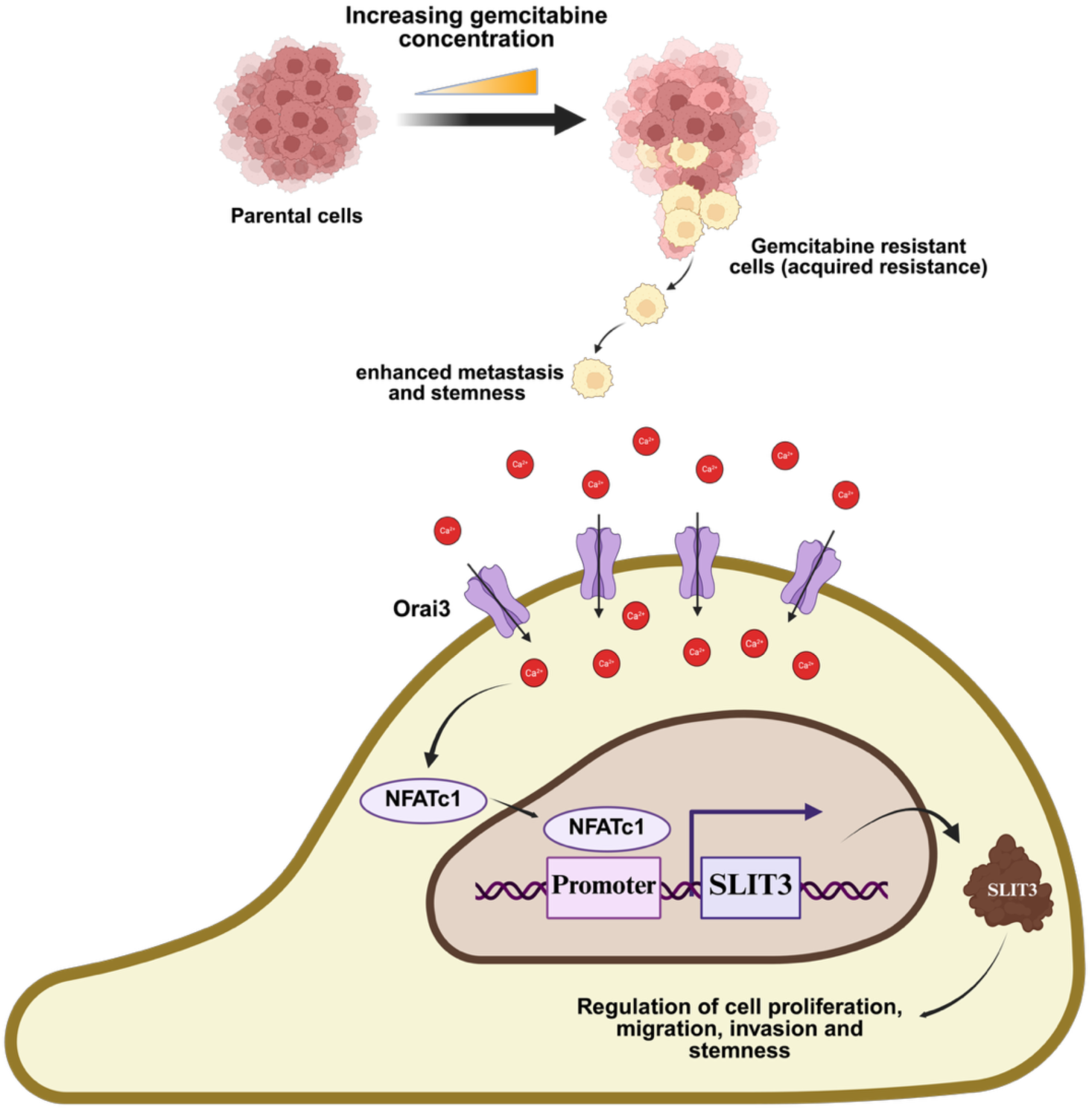

**Highlights:**

⇒ Orai3 is upregulated in gemcitabine-resistant pancreatic cancer cells
⇒ Orai3 is essential for chemo-sensitivity, migration, invasion and stemness *in vitro*
⇒ Orai3 regulates metastasis of gemcitabine-resistant pancreatic cancer cells *in vivo*
⇒ Unbiased RNA-seq reveals that SLIT3 functions downstream of Orai3 to drive gemcitabine resistance
⇒ NFATc1 transcription factor bridges Orai3 to SLIT3 transcription

## Introduction

Pancreatic Cancer (PC) is one of the leading causes of cancer-associated deaths worldwide, with a five-year survival rate of below 12% (Bray et al., 2024). PC is likely to surpass breast, prostate, and colorectal cancers to become the second leading cause of cancer-related death by 2040 (Rahib et al., 2014).The primary reasons for high mortality in PC is metastasis and chemoresistance. Currently, gemcitabine is the first-line therapy for treating PC. It is a deoxycytidine analogue inhibiting DNA replication and preventing tumor growth (Burris et al., 1997). Despite being the cornerstone of PC treatment for the last two decades, patients develop gemcitabine resistance within weeks of treatment initiation (Kim et al., 2008). Hence, there is an urgent need to overcome gemcitabine resistance by identifying the underlying mechanisms and targeting them to enhance gemcitabine efficacy (Vaibhav et al., 2024).

Emerging literature suggests that dysregulated Ca^2+^ signaling plays an important role in chemoresistance (Delisi et al., 2023; Fiorio Pla et al., 2016). Further, STIM1, an ER Ca^2+^ sensor, is reported to contribute to gemcitabine resistance in PC (Kutschat et al., 2021; Zhou et al., 2020). STIM1 senses endoplasmic reticulum (ER) Ca^2+^ and upon depletion of Ca^2+^ in the ER, it activates plasma membrane channels to bring in Ca^2+^ into the cytosol. Since ER Ca^2+^ stores regulate intracellular Ca^2+^ influx, this process is known as Store-operated calcium entry (SOCE). SOCE leads to increases in the cytosolic Ca^2+^ concentration required for several cellular processes, including cell proliferation, migration, differentiation, and apoptosis. Further, SOCE contributes to enhanced migration and invasion in cancer cells. Importantly, elevated expression of SOCE players is associated with poor prognosis in a variety of cancer types (Hammad et al., 2021; Shapovalov et al., 2021; Tanwar et al., 2020; Vashisht et al., 2015). Although there is some evidence supporting role of STIM1 in chemoresistance, the identity of Ca^2+^ influx channel responsible for driving gemcitabine resistance and the underlying molecular mechanism remains unappreciated. Moreover, there is limited information on targetable molecular players that contribute to acquired gemcitabine resistance, which presents a significant challenge for patients undergoing chemotherapy.

We here reveal that Orai3 Ca^2+^ channel plays a critical role in gemcitabine resistance via SLIT3 signaling. We generated stable gemcitabine-resistant cell lines derived from their sensitive parental line by subjecting them to progressive gemcitabine exposure. We analyzed the expression of key Ca^2+^ handling proteins in them and found elevated Orai3 and STIM1 levels.

To determine the role of Orai3 in chemoresistance, we conducted live-cell Ca^2+^ imaging, a variety of cellular assays and *in vivo* studies. These studies demonstrated that Orai3 drives gemcitabine resistance and contributes to cancer cell stemness. Moreover, we uncovered Orai3’s role in regulating the metastatic potential of gemcitabine-resistant cells *in vivo* by performing live animal imaging in zebrafish model. Mechanistically, our unbiased transcriptomics revealed an interesting differential regulation of SLIT3 downstream of Orai3 in gemcitabine-resistant cells. We performed SLIT3 gain-of-function as well as loss-of-function studies and identified a crucial oncogenic role of SLIT3 in the progression of gemcitabine-resistant PC. Our data demonstrate that SLIT3 regulates cell proliferation, migration, invasion, and stemness of gemcitabine-resistant PC cells downstream of Orai3. Finally, we identified that NFATc1 transcription factor connects Orai3 to SLIT3 transcription. Taken together, this study advances our understanding of the molecular mechanisms driving chemoresistance in PC. Importantly, it identifies Orai3 as a promising therapeutic target to overcome gemcitabine resistance.

## Results

### Generation and characterization of gemcitabine-resistant cell lines

To delineate the role of Ca^2+^ signaling in the development of PC chemoresistance, we first established stable gemcitabine-resistant (GemR) pancreatic cancer cell lines i.e. MiaPaCa-2 (non-metastatic) GemR, Panc-1 (Invasive) GemR, and CFPAC-1 (metastatic) GemR, from the parental MiaPaCa-2, Panc-1, and CFPAC-1 cell lines by exposing them to increasing doses of gemcitabine, as shown in the schematic diagram **(Fig. 1A)**. Cells were considered GemR once the maximal inhibitory concentration was significantly higher in GemR cells than parental cells. The IC_50_ values of GemR cells showed an increase compared to parental cell lines (MiaPaCa-2: 4.92µM vs MiaPaCa-2 GemR: 19.19µM, Panc-1: 5.41µM vs Panc-1 GemR: 8.02µM, CFPAC-1: 1.91µM vs CFPAC-1 GemR: 36.65µM) with a highest increase observed in CFPAC-1 GemR cells compared to the other two cell lines **(Fig. 1B).**

**Figure 1:**
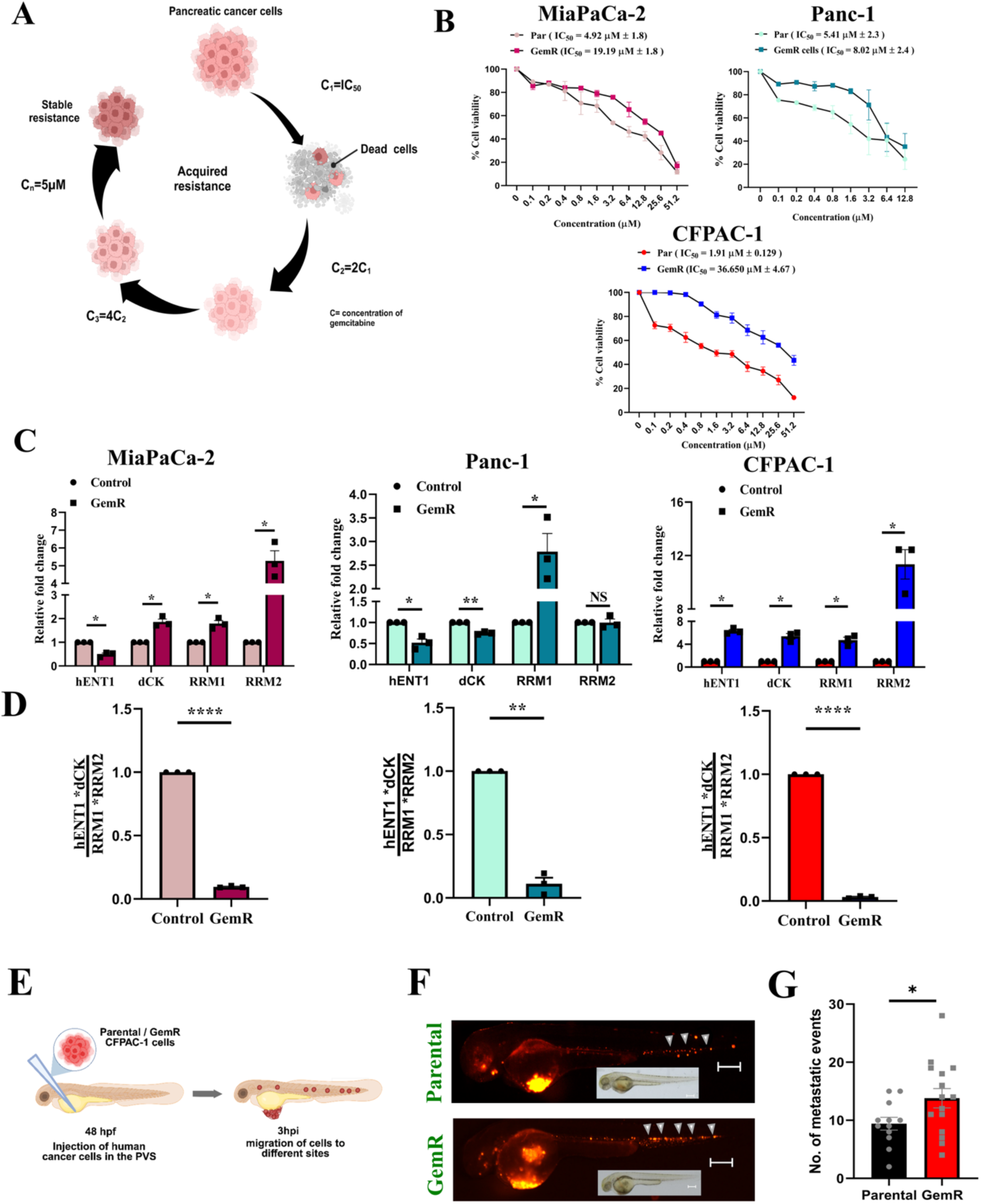
Generation and characterization of gemcitabine-resistant cells. **A)** Diagram illustrating the development of acquired gemcitabine-resistant cells. **B)** MTT assay showing IC50 values of Parental vs GemR in MiaPaCa-2, Panc-1 and CFPAC- 1 cell lines treated with gemcitabine for 72 h. The absorbance was normalized to the respective no-treatment absorbance (N=3). **C)** mRNA expression of *hENT1, dCK, RRM1,* and *RRM2* relative to *GAPDH* of GemR vs parental in MiaPaCa-2, Panc-1, and CFPAC-1 cell lines (N=3). **D)** Validation of gemcitabine resistance by *hENT1 x dCK / RRM1 x RRM2* expression ratio in GemR MiaPaCa-2, Panc-1, and CFPAC-1 cells. The value of the ratio in parental cells was set as 1.0 (N=3). **E)** Scheme representing the approach to inject pancreatic cancer cells (Parental, GemR, shNT, and shOrai3 GemR) in the Perivitelline Space (PVS) of 48 hpf *Assam wild type (Aswt)* zebrafish larvae. **F)** Representative images of zebrafish larvae 3 hours post-injection with parental and GemR labelled cells (red) captured using a stereomicroscope. Scale bar: 500 µm. **G)** Quantitative analysis of the metastatic capacity of CFPAC-1 parental and GemR cells upon engraftment into zebrafish (n=15). Data presented are mean ± S.E.M. For statistical analysis, a one-sample t-test was performed for panels C and D, and an unpaired test was performed for panel F using GraphPad Prism software. Here, * *p* <0.05; ** *p* < 0.01; *** *p* < 0.001 and **** *p* < 0.0001.

The transporters and enzymes that metabolize gemcitabine drive acquired GemR in human pancreatic cancer. Therefore, they are regarded as a predictive marker for response to gemcitabine in clinical settings. To validate the acquired resistance in our GemR cell lines, we quantified the expression of such well-established markers of GemR (Kim et al., 2008; Nakano et al., 2007) i.e. human equilibrative nucleoside transporter-1 (*hENT1*), deoxycytidine Kinase (*dCK*), Ribonucleotide reductase 1 (*RRM1*) and Ribonucleotide reductase 2 (*RRM2*). Our investigation showed a significant differential expression of these markers in the three gemcitabine resistance cell lines **(Fig. 1C)**. Further, as reported earlier, the acquired resistance in cell lines was estimated using the ratio of *hENT1* dCK / RRM1* RRM2* (Nakano et al., 2007). Our findings demonstrated higher IC_50_ and more pronounced acquired resistance in the metastatic CFPAC-1 compared to the two non-metastatic cell lines **(Fig. 1D)**. Therefore, we selected the CFPAC-1 cell line for the subsequent studies.

GemR pancreatic cancer is more aggressive, and it is associated with higher metastatic potential (Samulitis et al., 2015; Shah et al., 2007). Therefore, to assess the metastatic aggressiveness associated with GemR, we performed *in vivo* xenograft studies using zebrafish (*Danio rerio*) as a model system. As shown in the schematic diagram, we injected parental and GemR CFPAC-1 cells in the perivitelline space of 48 hours post-fertilization (hpf) old larvae. As reported earlier, the migration of cells at different sites post-injection was used to evaluate the metastatic potential of these cells (Stuelten et al., 2018; White et al., 2013). We observed that CFPAC-1 GemR cells showed aggressive metastatic behavior compared to the parental cells **(Fig. 1F,G)**. This observation confirmed the successful generation of a GemR cell line model with an enhanced metastatic ability as reported in pancreatic cancer patients.

### Orai3 expression and function is augmented in GemR cells

Earlier studies have shown a strong correlation between the expression of STIM proteins and chemoresistance (Kischel et al., 2019). We earlier reported that Orai3 encodes a functional SOCE channel in PCcells (Arora et al., 2021). However, role of Orai proteins in acquired PC chemoresistance remains unknown. Therefore, we first unbiasedly examined the expression of Orai and STIM proteins in parental and GemR cells. Interestingly, only Orai3 **(Fig. 2A,B)** and STIM1 **(Fig. 2C,D)** showed a significant upregulation in their expression. In contrast, other paralogs showed either a decrease or no change in their expression in GemR cells **(Fig. 2E and Supplementary Fig. 1)**, suggesting that Orai3 is uniquely overexpressed in GemR cells. Next, to investigate Orai3 activity in GemR cells, we performed live-cell Ca^2+^ imaging using Fura-2 AM. As reported in our previous study (Arora et al., 2021), SOCE was assessed using a standard thapsigargin (Tg) based protocol and 2APB treatment was given to distinguish between Orai1 and Orai3. Upon Tg-mediated ER Ca^2+^ store depletion, SOCE was studied by adding external Ca^2+^. Notably, we observed a significant increase in SOCE in GemR cells compared to parental cells **(Fig. 2F,G)**. Further, Ca^2+^ entry was potentiated by adding 2APB (50µM) and the 2APB potentiation was higher in GemR cells than parental cells **(Fig. 2G,H)**. This Ca^2+^ imaging data corroborates the protein expression data, thereby suggesting that GemR leads to SOCE augmentation by enhancing Orai3 and STIM1 expression.

**Figure 2:**
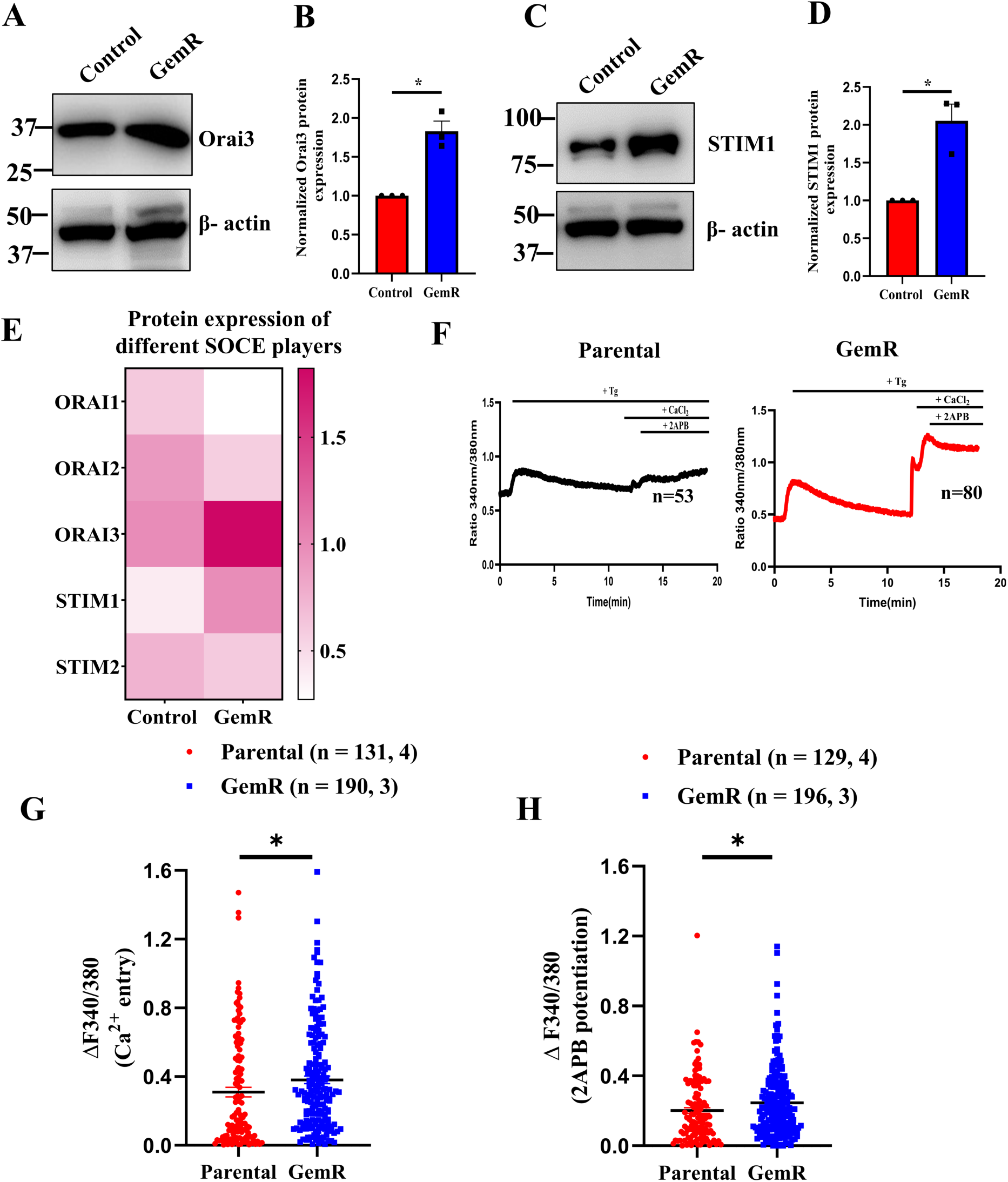
Orai3 expression and function is augmented in GemR cells. **A)** Representative western blot showing Orai3 expression in Parental and GemR CFPAC-1 cells. (N=3). **B)** Densitometric quantitation of Orai3 expression normalized to loading control β – actin. **C)** Representative western blot showing STIM1 expression in Parental and GemR CFPAC-1 cells (N=3). **D)** Densitometric quantitation of STIM1 expression normalized to loading control β – actin. **E)** Heatmap showing expression of different SOCE proteins in parental and GemR CFPAC-1 cells (N=3). **F)** Representative Ca^2+^ imaging traces of Parental and GemR CFPAC-1 cells where “n” denotes the number of cells in the trace. **G)** Quantitation of Ca^2+^ entry in Parental and GemR CFPAC-1 cells where “n=x,y” where “x” denotes the number of ROI in each condition and “y” denotes the number of dishes recorded. **H)** Quantitation of 2APB potentiation in Parental and GemR CFPAC-1 cells where “n=x,y” where “x” denotes the number of ROI in each condition and “y” denotes the number of dishes recorded. Data presented are mean ± S.E.M. For statistical analysis, a one-sample t-test was performed for panels B and D, while an unpaired student’s t-test was used for panels G and H using GraphPad Prism software. Here, * *p* <0.05; ** *p* < 0.01; *** *p* < 0.001 and **** *p* < 0.0001.

### Orai3 drives gemcitabine resistance *in vitro*

To investigate if Orai3 overexpression contributes to acquired chemoresistance, we generated Orai3 stable knockdown cell line in the background of GemR CFPAC-1 cells. Orai3 knockdown in the stable line was validated by performing quantitative RT-PCR. We observed a significant reduction in Orai3 expression in shOrai3 (shRNA specifically targeting Orai3) stable line compared to control shNT (non-targeting shRNA) line in GemR background **(Fig. 3A)**. Next, we investigated the proliferation ability of the four cell lines: parental, GemR, shNT GemR, and shOrai3 GemR. We observed that the proliferation rate in GemR and shNT GemR was higher than parental and shOrai3 GemR cells **(Fig. 3B)**. Interestingly, the increase in cell proliferation observed in GemR compared to parental cells was completely abrogated in shOrai3 GemR while it was comparable in shNT GemR **(Fig. 3B)**. This suggests that GemR cells have higher proliferative capabilities compared to parental cells, and Orai3 knockdown drastically decreases the enhanced proliferative capabilities of GemR cells. Next, we analyzed cell cycle progression using propidium iodide staining to investigate the underlying cause of the differential proliferation rates observed under these conditions. Our analysis revealed that both parental and shOrai3 GemR cells exhibited arrest at the G1/S phase. In contrast, GemR and shNT GemR cells showed significant progression into the S-phase, as indicated by the distribution of cell populations across different cell cycle stages **(Fig. 3C)**. The representative graphs of the cell cycle for all the conditions are shown (**Supplementary Fig. 2)**. Together, these findings demonstrate that Orai3 silencing significantly reduces proliferation of GemR cells, primarily due to cell cycle arrest in the G1/S phase.

**Figure 3:**
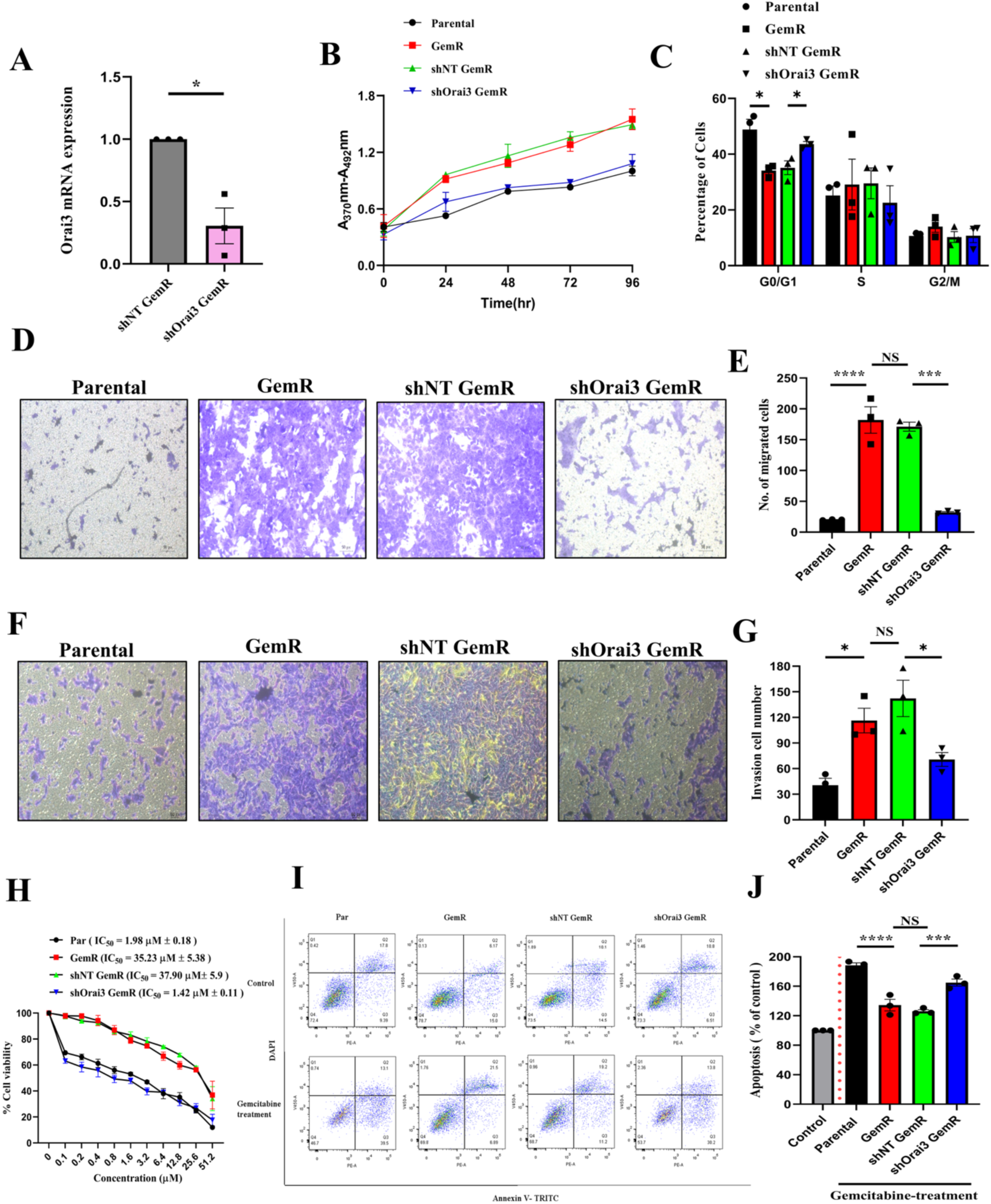
Orai3 regulates gemcitabine resistance *in vitro*. **A)** Knockdown validation of Orai3 lentiviral stables generated in CFPAC-1 GemR cells at qRT-PCR (N=3). **B)** BrdU proliferation assay representing proliferation rate upon silencing Orai3 at different time points. (N=3). **C)** Quantitation of cell number in different cell cycle phases calculated for four conditions (N=3). **D)** Boyden chamber transwell assay was performed to examine migration potential. Images were captured using a Nikon bright field microscope at 10x. **E)** The quantification of the number of cells migrated at the bottom of the transwell due to the FBS gradient is shown (N=3). **F)** Transwell assay for measuring invasion potential. Images were captured using a Nikon bright field microscope at 10x. **G)** The quantification of the number of cells that invade the matrigel coating on the transwell due to the FBS gradient is shown graphically (N=3). **H)** MTT assay demonstrating the alteration in gemcitabine sensitivity following Orai3 silencing in GemR CFPAC-1 cells, as reflected by changes in IC50 values (N=3). **I)** Flow cytometric apoptosis analysis performed using Annexin V and DAPI for parental, GemR, shNT and shOrai3 GemR CFPAC-1 cells alone or upon treatment with gemcitabine (N=3). **J)** Quantification of apoptotic cells under each condition is presented. Apoptotic cell counts were determined by summing early and late apoptotic cells and normalizing the total against their respective control cells. Data presented are mean ± S.E.M. One sample t-test was performed for panel A, an unpaired student t-test was performed for panel C using GraphPad Prism software, and a one-way ANOVA test was done for panels E, G, and J. Here, * *p* <0.05; ** *p* < 0.01; *** *p* < 0.001 and **** *p* < 0.0001.

Chemoresistance augments tumor cell migration and metastatic capabilities. Therefore, we studied the role of Orai3 in regulating migration and invasion potential of GemR cells. We performed transwell assays to assess the migratory and invasion potential of parental, GemR, shNT GemR and shOrai3 GemR CFPAC-1 cells. Our data revealed that migratory and invasion is highest in case of GemR and shNT GemR cells. In contrast, parental and Orai3-silenced GemR cells exhibited significantly reduced migration **(Fig. 3D,E)** and invasion **(Fig. 3F,G)**, This data demonstrates a positive correlation between Orai3 expression and migratory/invasion potential of GemR cells. Cell migration is a highly polarized process requiring actin cytoskeleton remodeling to coordinate protrusion, adhesion, contraction, and retraction. To examine the role of Orai3 in regulating the actin cytoskeleton in GemR cells, we performed phalloidin staining in shNT GemR and shOrai3 GemR CFPAC-1 cells. Our data revealed a significant shift in the distribution of actin filaments across cells under these conditions **(Supplementary Fig. 3A,B)**. Orai3 knockdown GemR cells showed actin distribution more at the center and less at the edges in contrast to shNT GemR cells, where the actin filaments are distributed more at the edges with a reduced presence towards the center. Therefore, our data suggests that Orai3 regulates actin dynamics of GemR cells, which in turn promotes the migration of these cells.

### Orai3 silencing sensitizes PC cells to gemcitabine by enhancing the rate of apoptosis

Next, we evaluated the role of Orai3 in protection against cytotoxic effects of gemcitabine. We silenced Orai3 in GemR PC cells and studied cytotoxicity of increasing doses of gemcitabine using the MTT cell viability assay. Our findings showed that Orai3 knockdown in GemR cells restored the sensitivity of gemcitabine-resistant cells, as reflected by a decrease in their IC_50_ values **(Fig. 3H)**. Interestingly, Orai3 silencing results in gemcitabine sensitivity similar to seen in case of parental cells. The heightened chemosensitivity to gemcitabine observed in shOrai3 GemR cells could be due to increased apoptosis. To investigate this, we performed apoptosis assays using flow cytometry. The four conditions of CFPAC-1 cells were treated with 5µM of gemcitabine for 72 h and stained with DAPI plus annexin V-TRITC labelled to distinguish between the apoptotic and necrotic cells. Our data shows a significant increase in apoptosis in parental and Orai3-silenced cells compared to GemR and shNT GemR cells **(Fig. 3I,J)**. Collectively, our data suggest that Orai3 is critical for providing a survival advantage to GemR cells by evading the process of apoptosis.

### Orai3 enhances the stemness of gemcitabine-resistant cells

Cancer stemness is a critical determinant of chemoresistance, contributing to recurrence and drug resistance in pancreatic cancer (Chaudhary et al., 2017; Zhang et al., 2016). To investigate Orai3’s contribution to the stemness of gemcitabine-resistant cells, we generated lentiviral-based stable knockdown of Orai3 in GemR cells. The stemness property of these cells was first confirmed through the 3-D spheroid formation assays. The size of the spheroid was used as a parameter to measure differences among four conditions. Interestingly, the bright-field image showed a well-formed and significantly larger spheroid in the case of GemR cells than parental, while Orai3-silenced cells generated spheroids smaller than shNT GemR cells **(Fig. 4A,B).** The GFP spheroid images of shNT and shOrai3 GemR are also shown. A seminal study had identified a subset of pancreatic cancer stem cells *in vivo*, which were CD44+ CD24+ ESA+ and harbored higher tumorigenic potential (C. Li et al., 2007). These triple-positive cells are enriched in gemcitabine-resistant pancreatic cancer cells(Shah et al., 2007). Further, B-cell-specific Moloney murine leukaemia virus insertion site 1 (BMI1) plays a crucial role in cancer pathogenesis. Pancreatic cancer stem cells isolated from both tumor spheres and xenografts based on the expression of CD44, CD24 and ESA, were shown to have significantly higher expression of BMI1 (Proctor et al., 2013). Therefore, we evaluated the expression of these key stemness players and observed a significant reduction in their expression upon silencing Orai3 **(Fig. 4C)**. Further, literature suggests that Aldehyde dehydrogenase (ALDH), a critical enzyme in regulating the retinoic acid pathway, and its isoforms are overexpressed in cancer stem cells (Ciccone et al., 2020; Deng et al., 2010; Marcato et al., 2011; Yue et al., 2022). In the context of pancreatic cancer, ALDH1A1 knockdown significantly reduced gemcitabine resistance, highlighting an important role of ALDH1A1 in acquired resistance to gemcitabine (Duong et al., 2012). Hence, we examined ALDH1A1 expression and observed that it was significantly upregulated in CFPAC-1 GemR cells compared to the control **(Fig. 4D, E)**. Importantly, upon Orai3 silencing in GemR cells, the expression of ALDH1A1 is significantly reduced. Taken together, our data demonstrate that Orai3 regulates cancer stemness properties as well as expression of key drivers of cancer stemness.

**Figure 4:**
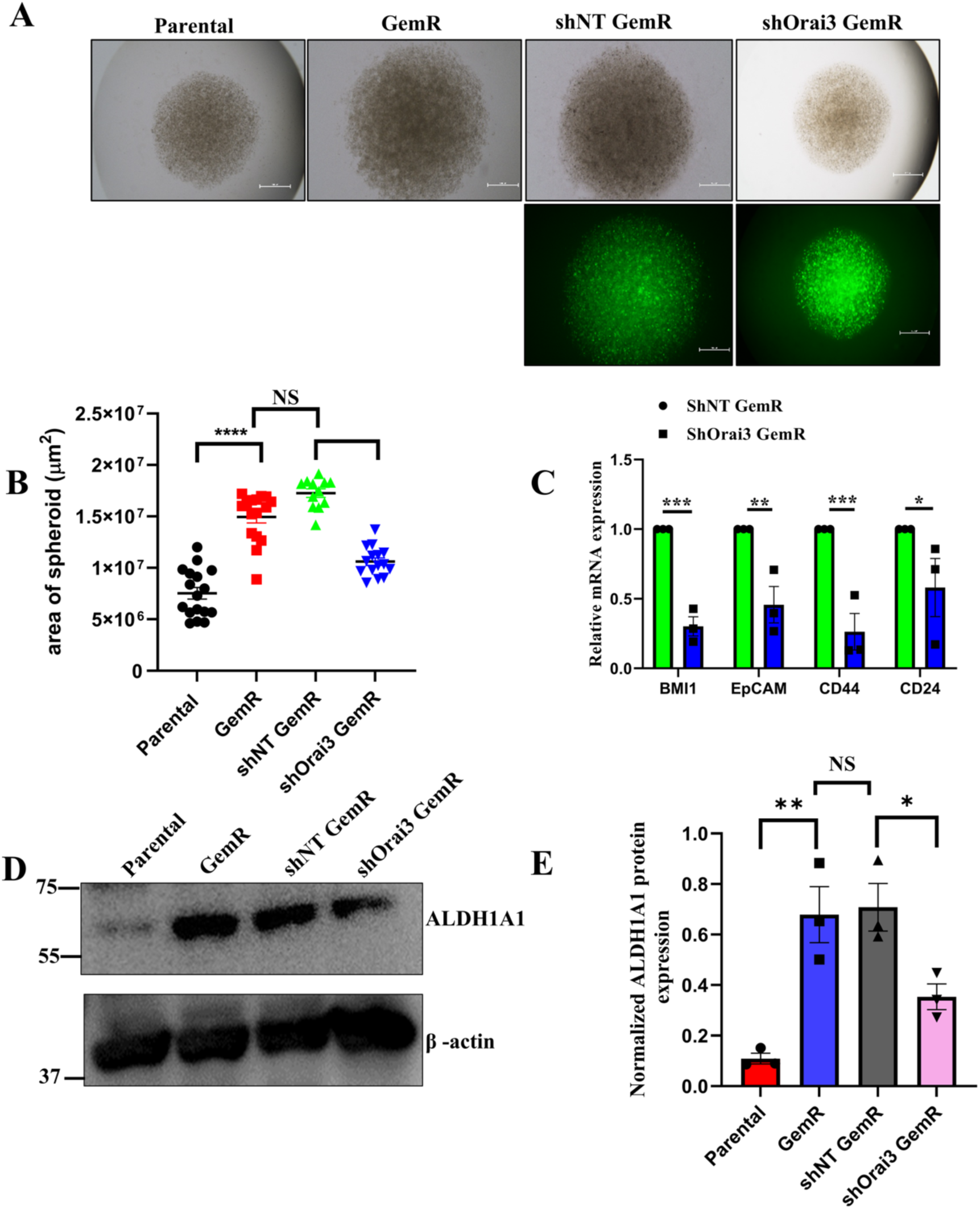
Orai3 regulates stemness of gemcitabine resistant cells. **A)** Spheroid formation assay conducted using the hanging drop method. **B)** Each spheroid was defined as a region of interest (ROI), and its area was measured using Nikon software. The quantified area is represented in the graph (N=3). **C)** qRT-PCR analysis demonstrates various stemness markers’ expression following Orai3 silencing in GemR CFPAC-1 cells (N=3). **D)** Representative western blot showing the expression of ALDH1A1 across different conditions. **E)** Densitometric analysis is done using ImageJ and presented graphically (N=3). The data presented is mean ± S.E.M. A one-way ANOVA test for panels B,E and a one-sample t-test was performed for panel C using GraphPad Prism software for statistical analysis. Here, * *p* <0.05; ** *p* < 0.01; *** *p* < 0.001 and **** *p* < 0.0001.

### Orai3 enhances metastatic potential of GemR cells *in vivo*

To study the role of Orai3 in regulating the metastatic potential of GemR cells *in vivo*, we used zebrafish (*Danio rerio*) as a model system. The initiation of the metastatic process involves migrating tumor cells from the primary tumor to other body parts to develop a secondary tumor. To replicate this process, we first optimized and generated CFPAC-1 xenografts in zebrafish. The four types of CFPAC-1 cell line i.e. parental, GemR, shNT GemR, and shOrai3 GemR, were labelled with CM-DiI (a red colored dye) and injected into the perivitelline space (PVS) of zebrafish embryos at 2 days post-fertilization (dpf). Subsequently, to determine the metastatic potential, each xenograft was analyzed at 4 days post-injection (dpi) to assess for micro-metastasis in the caudal hematopoietic tissue (CHT). The representative images of Parental, GemR, shNT, and shOrai3 GemR xenografts with metastasis in CHT are shown (**Fig. 5A**), and the percentage of metastasis is calculated using the following formula.

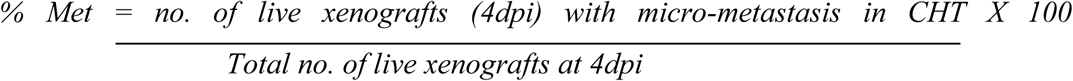

**Figure 5:**
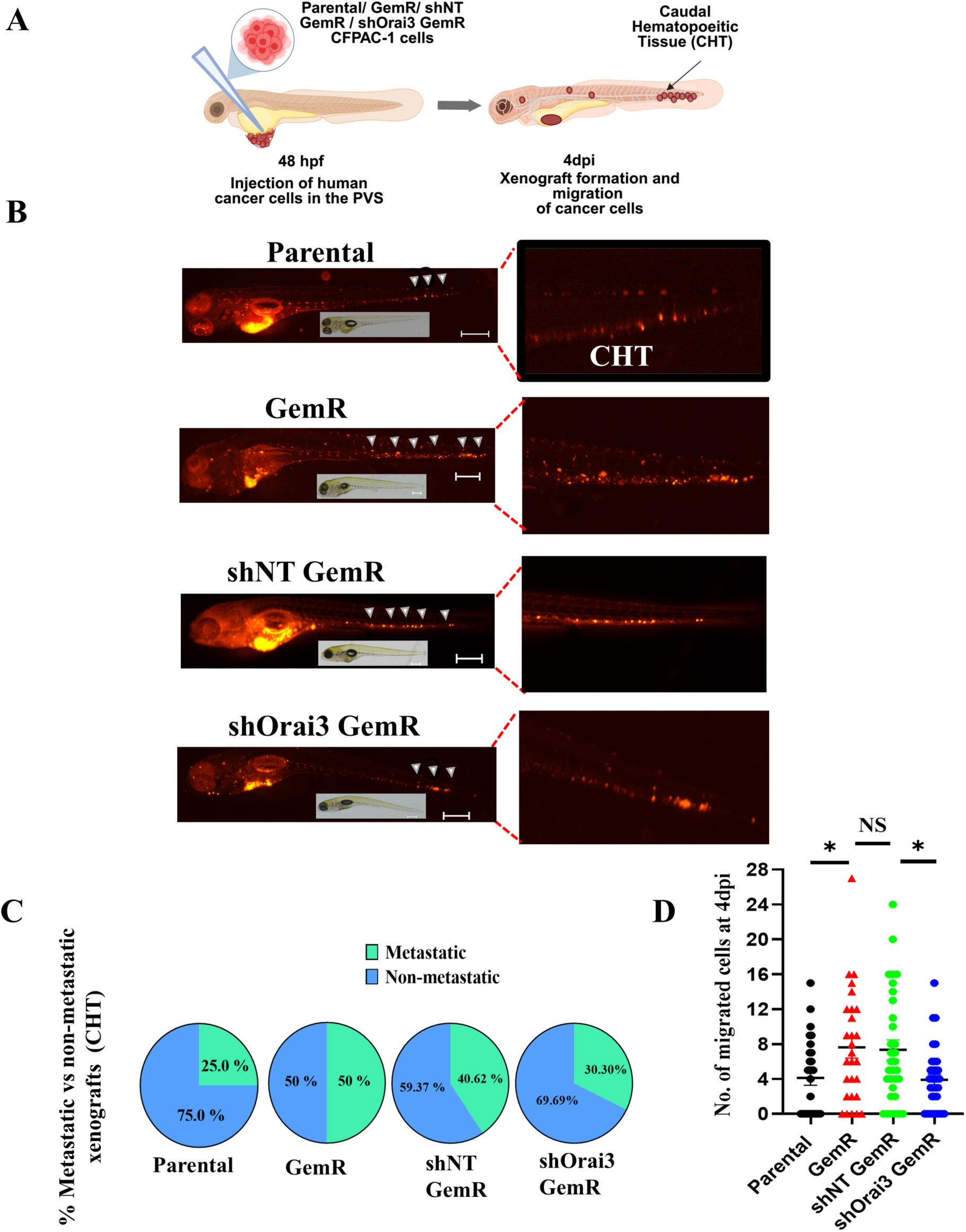
Orai3 regulation of GemR metastatic potential *in vivo*. **A)** Representative zebrafish xenograft images showing migration of labeled cells injected in PVS at 2dpf to CHT at 4dpi. **B)** The percentages of metastatic vs non-metastatic xenografts in different conditions are shown in the pie chart. Parental (n=28), GemR (n=28), shNT GemR (n=32), and shOrai3 GemR (n=33). **C)** Quantitation of the total number of cells migrated to different locations in xenografts is shown graphically. Data presented are mean ± S.E.M. A one-way ANOVA was performed for panel D using GraphPad Prism software. Here, * *p* <0.05; ** *p* < 0.01; *** *p* < 0.001 and **** *p* < 0.0001. The ‘n’ denotes the total number of xenografts analyzed for quantitation.

Using this formula, we calculated that 50.0% and 40.62% xenografts in GemR and shNT GemR cells exhibited metastasis to the CHT, while Orai3 silencing reduced this to 30.30%, which is comparable to parental cell xenografts that showed 25.0% metastatic xenografts **(Fig. 5B)**. Additionally, we quantified the number of cells that metastasize to different sites in the zebrafish body at 4dpi using ImageJ software. As expected, our data revealed that GemR cells have higher metastatic ability compared to parental cells. Importantly, we observed a significant reduction in metastatic cells in shOrai3 GemR as compared to shNT GemR cells **(Fig. 5C)**. Taken together, our findings highlight a crucial role for Orai3 in driving the metastasis of GemR cells *in vivo*.

### Unbiased transcriptomics reveals that SLIT3 functions downstream of Orai3

Our *in vitro* and *in vivo* data demonstrate that Orai3 plays a pivotal role in sustaining gemcitabine resistance in pancreatic cancer cells. Next, we performed bulk RNA seq to unveil the molecular mechanism that contribute to acquired gemcitabine resistance downstream of Orai3. RNA seq was performed on all four conditions: parental, GemR, shNT GemR and shOrai3 GemR CFPAC-1 cells **(Fig. 6A)**. The fold change was calculated for test conditions GemR and shOrai3 GemR with respect to (w.r.t.) their respective control, i.e., parental and shNT GemR. Genes showing a log2 fold change of +2 w.r.t. control were considered upregulated genes, while a cut-off log2 fold change of less than -2 was set to filter out downregulated genes. The differentially expressed genes (DEGs) were screened according to |logFC| > 2 and P ≤0.05. The differentially expressed genes for GemR vs parental **(Supplementary Fig. 4A)** and shOrai3 GemR vs shNT GemR **(Supplementary Fig. 4B)** groups are shown in the volcano plots. We also examined the biological, cellular, and molecular pathways associated with gemcitabine resistance through GO pathway enrichment analysis in the group GemR vs parental **(Supplementary Fig. 4C**,D) and shOrai3 GemR vs shNT GemR **(Supplementary Fig. 4E**,F). To identify the genes that are common but differentially regulated in parental vs GemR and shNT vs shOrai3 GemR groups, we looked for common genes which were upregulated in the GemR cells vs parental but downregulated in shOrai3 GemR in comparison to shNT GemR. This analysis led to the identification of 12 genes that showed a differential expression profile. Since the Orai3-mediated effect would be most likely due to its Ca^2+^ signaling function, we further overlapped these 12 genes with the Ca^2+^ associated genes database (Hörtenhuber et al., 2017). This database comprises all the genes implicated in Ca^2+^ signaling and are linked to human disorders. This robust analysis revealed only one gene, SLIT3, which was common among the three groups **(Fig. 6B)**. SLIT3 belongs to a family of secretory proteins that bind to the roundabout guidance receptor (ROBO) family of transmembrane proteins, serving as an axonal guidance cue in developing the nervous system. (Yuan et al., 1999). Further, recent studies have shown that SLIT3 is crucial for cancer initiation and progression (Mehlen et al., 2011).

**Figure 6:**
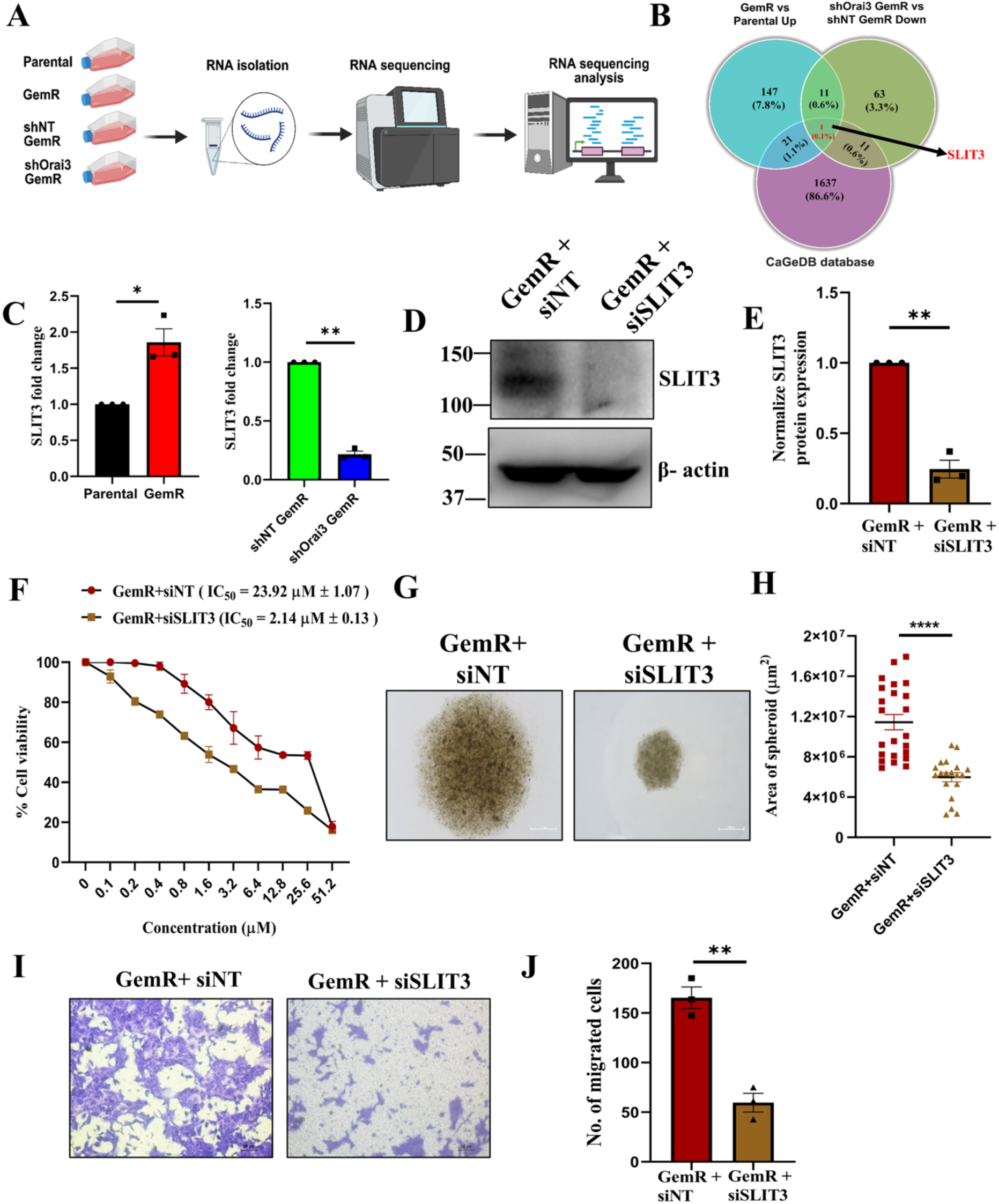
SLIT3 positively regulates gemcitabine resistance. **A)** Schematic diagram showing steps for RNA-sequencing workflow. **B)** Venn diagram representing common differentially expressed genes in GemR vs Parental up overlapped and shOrai3 GemR vs shNT GemR down overlapped with Ca^2+^-regulated genes. **C)** Validation of SLIT3 mRNA expression in parental vs GemR and shNT vs shOrai3 GemR CFPAC-1 cells (N=3). **D)** Representative western blot confirming siRNA-based silencing of SLIT3 in GemR cells. **E)** Densitometric analysis of SLIT3 knockdown using imageJ is shown graphically (N=3). **F)** MTT assay shows the change in the chemosensitivity of GemR cells upon silencing SLIT3 (N=3). **G)** Spheroid formation assay depicting a spheroid’s size change on silencing SLIT3 in GemR cells. Each spheroid was defined as a region of interest (ROI), and its area was measured using Nikon software. **H)** The quantified area of spheroids is represented in the graph (N=3). **I)** Boyden chamber transwell assay for studying migration potential of GemR cells upon silencing SLIT3 (N=3). **J)** Graph showing quantitative analysis of migration potential in SLIT3 knockdown cells. Data presented are mean ± S.E.M. A one-sample t-test was performed for panels C and E and an unpaired student t-test for panels H and J using GraphPad Prism software. Here, * *p* <0.05; ** *p* < 0.01; *** *p* < 0.001 and **** *p* < 0.0001.

Interestingly, whole-genome studies on PC patient samples have revealed mutations in the SLIT and ROBO genes (Aung et al., 2018; Biankin et al., 2012; Waddell et al., 2015). These studies have highlighted a strong correlation between the dysregulated SLIT/ROBO signaling and early stages of pancreatic neoplastic transformation (Biankin et al., 2012). However SLIT3’s role in pancreatic cancer chemoresistance remains unappreciated. Therefore, based on the transcriptomics data analysis and literature survey, we focused on understanding the role of SLIT3 in Orai3-induced chemoresistance. We first examined the expression of SLIT3 in the samples used for RNA sequencing to validate our transcriptomics results. As observed in transcriptomics, SLIT3 expression is higher in GemR cells in comparison to parental cells, while it decreases in the Orai3-silenced GemR cells as compared to shNT control GemR cells **(Fig. 6C)**. To understand the role of SLIT3 in gemcitabine resistance, we started by silencing SLIT3 in GemR cells using siRNA. As shown in our western blotting data, there is a robust reduction in SLIT3 protein levels upon transfection with siRNA targeting SLIT3 **(Fig. 6D,E)**. subsequently, the siSLIT3 transfected cells were used to study the role of SLIT3 in chemoresistance. First, we performed MTT assays to examine the role of SLIT3 in chemosensitivity. We observed that SLIT3 knockdown in these cells increases sensitivity to gemcitabine, as we see a robust (10-fold) decrease in IC_50_ values of gemcitabine **(Fig. 6F)**. To examine the effect of SLIT3 silencing on stemness, we performed 3-D spheroid assays that revealed a drastic reduction in the spheroid area **(Fig. 6G,H)**. Furthermore, the transwell migration studies show that SLIT3 knockdown in GemR cells results in a significant decrease in migration potential compared to control **(Fig. 6I,J)**. Altogether, our results uncover that downstream of Orai3, SLIT3 expression is augmented in GemR cells. Further, SLIT3 plays an important role in enhancing chemoresistance, migration and stemness of GemR cells thereby driving gemcitabine resistance.

### SLIT3 drives gemcitabine resistance downstream of Orai3

We observed that SLIT3 expression is increased in GemR cells, and it is decreased in Orai3- silenced GemR cells. Further, we found that SLIT3 contributes to GemR in pancreatic cancer cells. Next, to examine whether SLIT3 contributes to gemcitabine resistance downstream of Orai3, we performed overexpression of SLIT3 in Orai3-silenced GemR cells. SLIT3 was transfected in Orai3-silenced GemR cells, resulting in significant overexpression of SLIT3 **(Fig. 7A,B)**. We subsequently examined chemosensitivity of these cells by performing MTT assays at different gemcitabine concentrations. These experiments demonstrated a substantial increase in chemoresistance in SLIT3 overexpressing Orai3 silenced GemR cells compared to the Orai3 silenced GemR cells **(Fig. 7C)**. Further, 3-D spheroid formation assays showed enhanced stemness properties **(Fig. 7D,E)** and transwell assays uncovered higher migration **(Fig. 7F,G)** & invasion **(Fig. 7H,I)** potential in SLIT3 complemented cells in comparison to Orai3 silenced GemR cells. Collectively, our data reveal that overexpression of SLIT3 in Orai3-silenced GemR cells restores chemoresistance, migration, invasion and stemness. Hence, this data confirms that SLIT3 works downstream of Orai3 to drive gemcitabine resistance.

**Figure 7:**
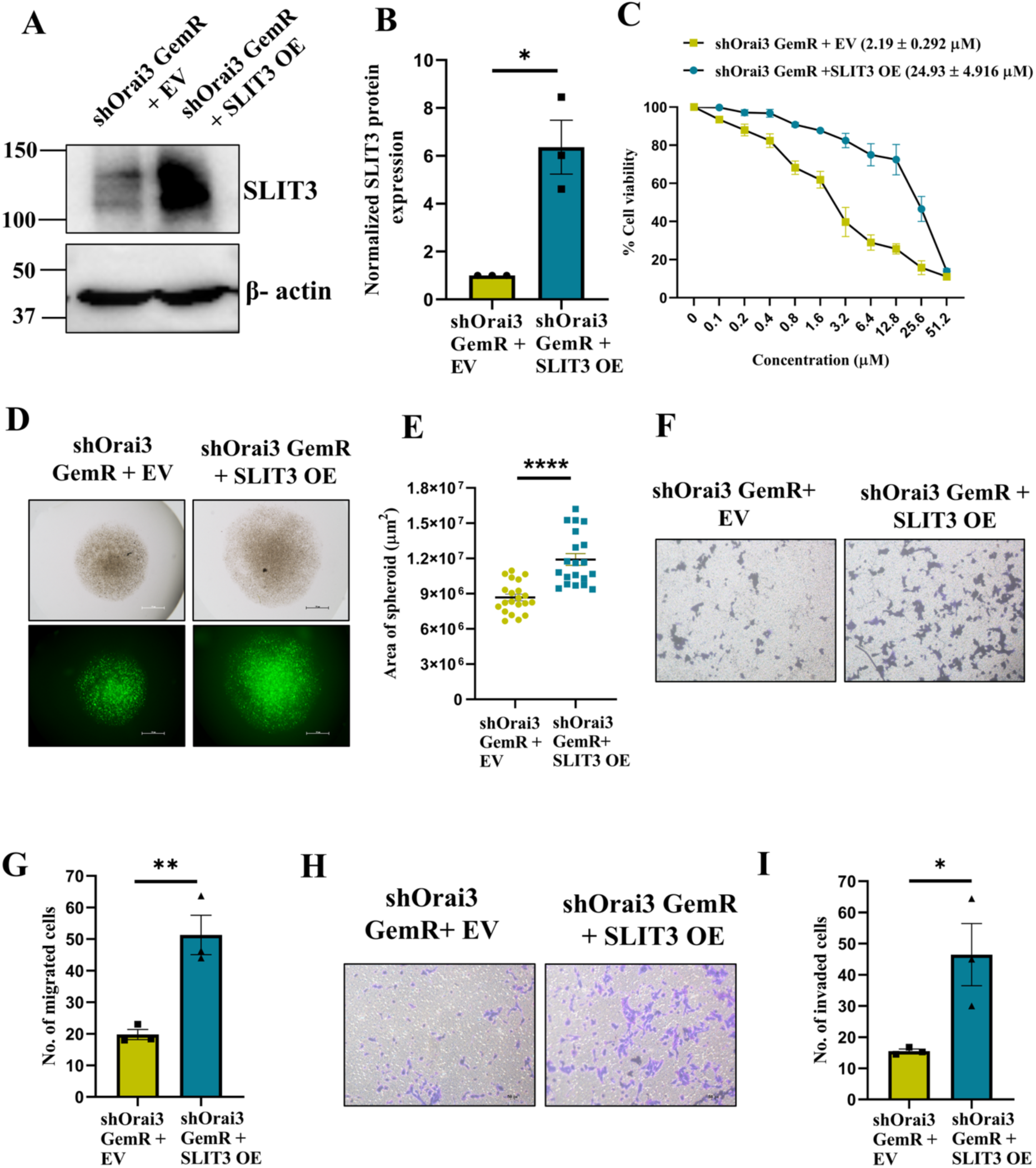
SLIT3 functions downstream of Orai3 to regulate gemcitabine resistance. **A)** Representative western blot confirming overexpression SLIT3 in Orai3-silenced GemR cells. **B)** Densitometric blot analysis using imageJ is shown graphically (N=3). **C)** MTT assay shows the change in chemosensitivity of shOrai3 GemR cells following overexpression of SLIT3 compared to empty vector control (N=3). **D)** Spheroid formation assay depicting the shape change of spheroid upon overexpression of SLIT3 in shOrai3 GemR cells. Each spheroid was defined as a region of interest (ROI), and its area was measured using Nikon software. **E)** The quantified area of spheroids is represented in the graph (N=3). **F)** Boyden chamber transwell assay showing migration potential of shOrai3 GemR cells upon overexpression of SLIT3. **G)** Quantitation of migration is shown graphically (N=3). **H)** Boyden chamber transwell assay with Matrigel coating showing invasion ability of shOrai3 GemR cells upon overexpression of SLIT3. **I)** Quantitation of invasion is shown graphically (N=3). Data presented are mean ± S.E.M. A one-sample t-test was performed for panels B and an unpaired student t-test for panels E, G, I using GraphPad Prism software. Here, * *p* <0.05; ** p < 0.01; *** *p* < 0.001 and **** *p* < 0.0001.

### SLIT3 expression is transcriptionally regulated by NFATc1

Our transcriptomics data identified SLIT3 as a differentially regulated gene downstream of Orai3 in GemR cells. To delineate the molecular mechanism driving SLIT3’s transcriptional regulation by Orai3, we investigated potential role of the Nuclear Factor of Activated T cells (NFAT) transcription factors. The activation of NFATc1-c4 requires a sustained Ca^2+^ signal, which is achieved by the opening of Orai channels at the plasma membrane (Hogan et al., 2003, 2010). We performed extensive bioinformatic analysis to examine the potential binding sites of NFATs on SLIT3 promoter using two independent tools, namely the Eukaryotic Promoter Database (EPD) search motif tool (https://epd.expasy.org/epd/) and PSCAN (Tang et al., 2020). The databases revealed five potential binding sites for NFATc2 and four binding sites for NFATc1 and NFATc3 on the SLIT3 promoter after applying the stringent p-value cut off of 0.001 **(Fig. 8A)**, while no binding site for NFATc4 was predicted by these tools. PSCAN was used to identify the sequence of putative TF-binding sites present on the SLIT3 promoter **(Fig. 8B)**. This was done using the JASPAR Core 2020 non-redundant transcription factor position weight matrix database. These two tools, each using a different algorithm, identified putative binding sites and a putative consensus sequence present on the SLIT3 promoter for different NFATs. Thus, our robust bioinformatic analysis suggests that NFATs can potentially regulate SLIT3 transcription downstream of Orai3.

**Figure 8:**
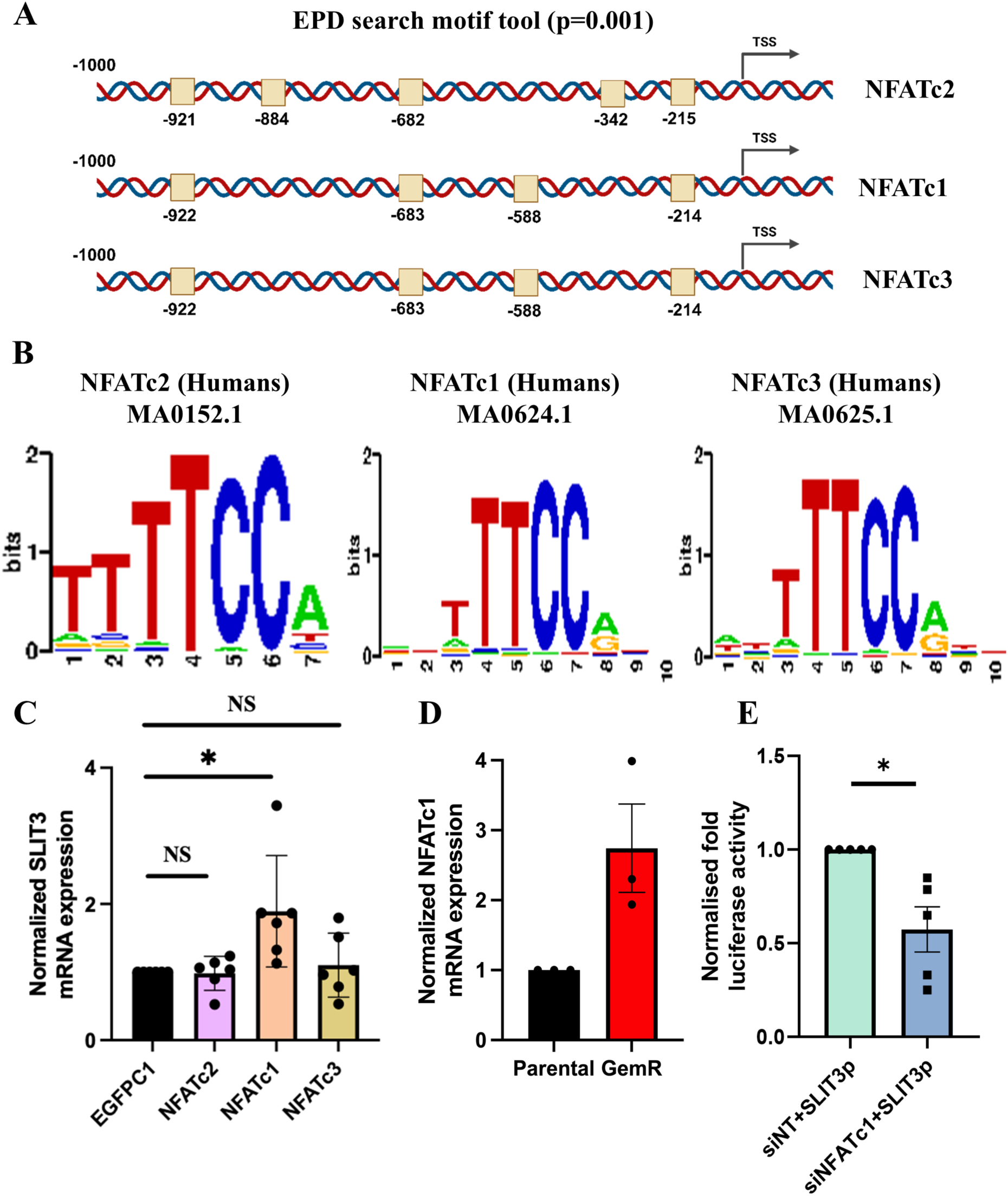
SLIT3 is transcriptionally regulated by NFATc1 in GemR cells. **A)** Identification of putative binding sites on SLIT3 promoter using EPD tool at a p-value cut-off of 0.001. **B)** A position weight matrix of human NFATc2/c1/c3 consensus binding sequence on SLIT3 promoter. **C)** SLIT3 mRNA expression following NFATc2/c1/c3 overexpression in the Parental CFPAC- 1 cell line (N=5). **D)** Normalised luciferase activity of SLIT3 promoter upon NFATc1 silencing in CFPAC-1 GemR cells (N=5). Data presented are mean ± S.E.M. A one-way ANOVA was performed for panel C and a one- sample t-test for panel D using GraphPad Prism software. Here, * *p* <0.05; ** *p* < 0.01; *** *p* < 0.001 and **** *p* < 0.0001.

To identify which NFAT isoform is regulating the expression of SLIT3, we performed overexpression studies in the wild-type CFPAC-1 cell line, in which SLIT3 expression is low. We observed that NFATc1 overexpression markedly increased SLIT3 mRNA levels compared to empty vector, while the expression of SLIT3 was unchanged upon NFATc2 or NFATc3 overexpression **(Fig. 8C)**. We further performed luciferase assays to assess the SLIT3 promoter activity in response to NFATc1 overexpression. Since NFATc1 expression is higher in CFPAC-1 GemR cells **(Fig. 8D)**, we silenced NFATc1 in them and simultaneously overexpressed SLIT3 promoter encompassing all four predicted SLIT3 binding sites cloned into the dual luciferase reporter vector, psi-CHECK 2. After 48 hrs of transfection, we performed luciferase assays to measure promoter activity. The results revealed a marked decrease in SLIT3 promoter activity following NFATc1 silencing **(Fig. 8E)**, strongly suggesting that NFATc1 positively regulates SLIT3 transcription and is a key driver of its activation in GemR cells. These findings further support our bioinformatic analysis and establish a functional role of NFATc1 in regulating SLIT3 transcription.

In summary, we reveal that Orai3 plays a crucial role in acquired gemcitabine resistance in PC. We performed a variety of *in vitro* assays and *in vivo* studies to demonstrate that Orai3 silencing enhances gemcitabine sensitivity in the resistant cells. Further, we performed unbiased transcriptomics analysis and identified that SLIT3 drives gemcitabine resistance downstream of Orai3. Moreover, we delineated that NFATc1 transcription factor connects Orai3 to SLIT3 regulation **(Fig. 9)**. In future, the strategies for targeting Orai3 in combination with gemcitabine may pave the path for more efficient chemotherapy regime against pancreatic cancer.

**Figure 9:**
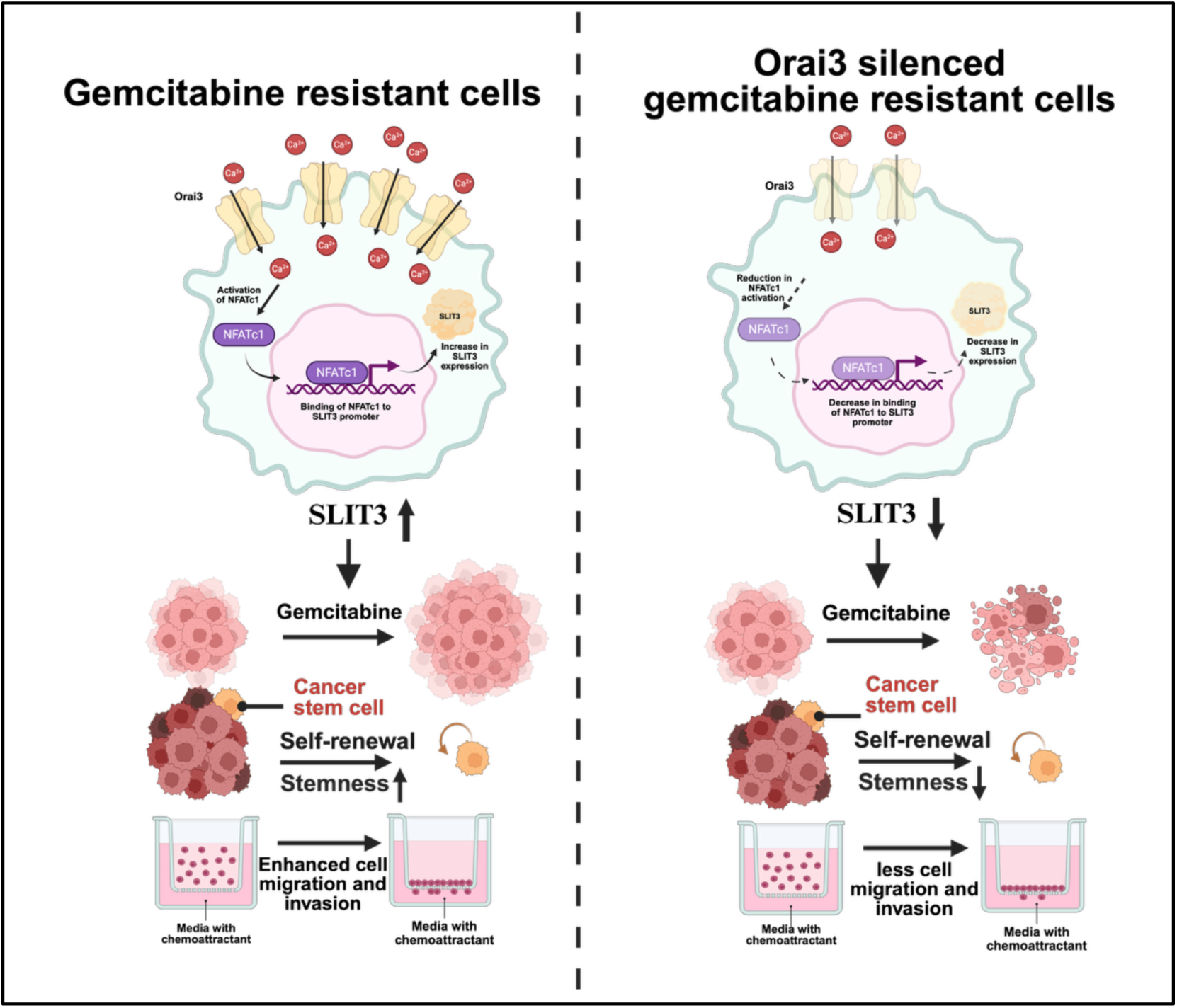
Orai3 drives pancreatic cancer gemcitabine resistance via NFATc1-SLIT3 axis. The graphical summary of our work demonstrates that Orai3 activates NFATc1, which in turn transcriptionally regulates the expression of SLIT3. The NFATc1-SLIT3 axis functions in coordination to regulate gemcitabine resistance downstream of Orai3. Silencing of Orai3 results in a reduction of NFATc1 activation, thereby affecting SLIT3 transcription, which in turn disrupts chemoresistance and its key hallmarks such as cell proliferation, migration, invasion, and stemness.

## Discussion

Pancreatic Cancer (PC) is one of the deadliest cancers and is associated with an extremely poor prognosis. Gemcitabine is the first line of chemotherapy used for the management of PC. However, acquired resistance to gemcitabine remains a major clinical challenge, significantly limiting gemcitabine efficacy. With the limited effective second-line treatment options available, there is a critical need to develop novel therapeutic strategies aimed at overcoming gemcitabine resistance.

Ca^2+^ signaling is emerging as a key driving force in mediating the cytotoxic effects of chemotherapy as numerous chemotherapeutic agents trigger an extensive rewiring of cytosolic Ca^2+^ signaling in cancer cells (Kischel et al., 2019). This phenomenon is primarily attributed to the dysregulated expression or activity of various plasma membrane Ca^2+^ ion channels (Bhatnagar et al., 2025; Vashisht et al., 2015). In non-excitable cells, Ca^2+^ entry is mainly driven through a process known as Store Operated Ca^2+^ Entry (SOCE) and is mediated by the family of Orai channels, namely Orai1, Orai2, and Orai3 (Tanwar et al., 2020). Evolving evidence indicates that enhanced SOCE plays a crucial role in the development of chemoresistance. A recent study showed co-amplification of RRM1 and STIM1 in gemcitabine-resistant pancreatic cancer cells. The heightened expression of STIM1 correlates with elevated store-operated Ca^2+^ entry (SOCE) and dysregulated endoplasmic reticulum (ER) stress, resulting in enhanced NFATc2 activity and epigenetic reprogramming (Kutschat et al., 2021). Another intriguing study developed a gemcitabine-resistant pancreatic cancer cell line and used high-throughput omics platforms to identify calmodulin as a key driver of gemcitabine resistance. Inhibiting calmodulin with a Ca^2+^ channel blocker, amlodipine restored the gemcitabine sensitivity *in vitro* and *in vivo* (Principe et al., 2022). Collectively, these studies highlight that augmented Ca^2+^ influx contributes to gemcitabine resistance, but the identity of the Ca^2+^ channel and downstream molecular mechanism remains elusive.

Orai3 encodes a SOCE channel in a sub-set of breast cancer cells and drives breast tumorigenesis (Motiani et al., 2010; Motiani, Zhang, et al., 2013; Vashisht et al., 2018). Further, our previous study demonstrated that Orai3 functions as a SOCE channel in PC and is essential for PC progression & metastasis (Arora et al., 2021). However, the functional relevance of Orai3 in acquired gemcitabine resistance remains unappreciated. Here, we report a novel role of Orai3 in acquired gemcitabine resistance. We show that gemcitabine-based chemotherapy selectively increases Orai3 and STIM1 expression. In contrast, the expression levels of Orai1, Orai2, and STIM2 remains unaffected in metastatic CFPAC-1 cells. The other two cell lines, i.e., MiaPaCa-2 and Panc-1, used in this study exhibit relatively lower sensitivity to gemcitabine, as reflected by the fold change in IC_50_ values between parental and GemR cells. This disparity likely reflects intrinsic differences in gemcitabine resistance among these cell lines. Indeed, previous studies have reported that in these cells gemcitabine IC_50_ concentrations ranged from tens to hundreds of nanomolar, making them less suitable to study gemcitabine resistance as compared to CFPAC-1 (Arumugam et al., 2009; Huanwen et al., 2009). Importantly, our data demonstrated higher IC_50_ and more pronounced acquired gemcitabine resistance in CFPAC-1 compared to MiaPaCa-2 and Panc-1. Therefore, we selected CFPAC- 1 cells for delineating the role of Orai3 in acquired gemcitabine resistance.

Another intriguing observation was the increase in Ca^2+^ entry in GemR cells, highlighting gemcitabine’s influence on cytosolic Ca^2+^ levels. The gemcitabine resistance augments SOCE **(Fig. 2F)** and upregulates Orai3 and STIM1 **(Fig. 2A-D)**. Furthermore, the enhanced 2-APB- mediated potentiation of Ca^2+^ entry in GemR cells confirms the functional role of Orai3 in these cells **(Fig. 2F, H)**. This data suggests that gemcitabine resistance results in elevation of Ca^2+^ entry that may impact drug sensitivity and may induce resistance mechanisms. Here, we reason that cancer cells may upregulate SOCE during chemotherapy as a survival mechanism. This adaptive response might help cancer cells to maintain Ca^2+^ homeostasis and activate pro-survival signals to resist chemotherapy-induced cell death. Therefore, targeting SOCE proteins under these conditions could disrupt the protective mechanism, making cancerous cells more vulnerable to chemotherapy. Indeed, our data clearly demonstrates that Orai3 plays a crucial role in acquired chemoresistance, and its silencing makes cells more susceptible to gemcitabine **(Fig. 3H).** This suggests that targeting Orai3 can be a potential therapeutic strategy to enhance the efficacy of gemcitabine by sensitizing resistant cells to therapy.

The role of Orai proteins in the induction of cancer stem cell (CSC) phenotype has been highlighted in various cancers such as glioblastoma (Liu et al., 2011; Motiani, Hyzinski-García, et al., 2013) and lung cancer (Daya et al., 2021; Hou et al., 2011; W. Li et al., 2013). The CSC properties include self-renewal ability, tumor spheroid formation, drug resistance, enhanced migratory potential, elevated expression of CSC markers (Beck et al., 2013). Gemcitabine resistance can augment the CSC subpopulation through different pathways, such as Nox/ROS/NF-κB/STAT3 and hypoxia-induced Akt/Notch1 signalling pathways (Zhang et al., 2016, 2018). In our study, we demonstrate that Orai3 silencing in GemR cells results in reduction in the expression of stemness markers CD44, CD24, EPCAM, BMI1 and ALDH1A1 **(Fig. 4C-E)**. Further, Orai3 plays a critical role in tumor spheroid formation **(Fig. 4A,B)** and contributes to enhanced metastasis of GemR cells **(Fig. 5A-C)**. Collectively, our data reveals that Orai3 is crucial for regulating the stemness of GemR PC cells.

Cancer cell migration is a crucial step in metastasis and is regulated by changes in the structure and dynamics of the actin cytoskeleton. It is reported that spatiotemporal variations in intracellular Ca^2+^ levels have a key role in forming new protrusions at the leading edge and disassembling mature adhesions at the rear (Lauffenburger et al., 1996). Our study showed that silencing Orai3 significantly reduced the migration potential of CFPAC-1 GemR cells **(Fig. 3D, E)**. To examine the underlying mechanism, we analyzed the distribution of actin filaments across the cell by staining them with phalloidin. Interestingly, we observed that in Orai3 silenced GemR cells, actin filaments were distributed more towards the nucleus and less at the periphery of cells than in control cells **(Supplementary Fig. 3A,B)**. Recently, the connection between actin remodeling and chemoresistance was reported, where it was shown that RhoJ facilitates EMT-associated resistance to chemotherapy (Debaugnies et al., 2023). Rho GTPases are a family of small G-proteins that control tumor initiation and progression by regulating cell migration, proliferation, apoptosis and by influencing metabolism, senescence and cancer cell stemness (Crosas-Molist et al., 2022). Analysis of our transcriptomics data showed downregulation of Rho GTPases in Orai3-silenced GemR cells compared to control **(Supplementary Fig. 3C)**. This suggests that Orai3 regulates the actin dynamics of GemR cells, thereby enhancing their migratory and metastatic potential.

For unbiased identification of the molecular mechanism working downstream of Orai3 to drive gemcitabine resistance, we performed RNA-seq analysis on parental, GemR, shNT and shOrai3 GemR cells. Our transcriptomics analysis identified SLIT3 **(Fig. 6B)**, which acts downstream of Orai3 to induce gemcitabine resistance. SLITs are a class of secretory glycoprotein that binds to the roundabout guidance receptor (ROBO) family of transmembrane proteins, serving as an axonal guidance cue in developing the nervous system. Interestingly, SLITs are also known to protect pancreatic beta-cells from ER-stress-induced apoptosis via the gradual release of Ca^2+^ from ER stores(Yang et al., 2013). Functionally, SLITs ligands along with ROBO receptors play an important role in attracting and repelling neuronal migration. This dual functionality is also observed in different cancers where SLITs can act both as a tumor suppressor and an oncogene. So far, the role of SLIT3 in PC progression and chemoresistance is unknown. Our study reveals that SLIT3 drives PC cell proliferation, stemness and migration (**Fig. 6F-J**). Further, our data demonstrates that SLIT3 acts downstream of Orai3 and positively regulates chemoresistance, stemness, migration and invasion of gemcitabine-resistant cells (**Fig. 7**). Therefore, our study identifies SLIT3 and its crosstalk with Orai3 as a crucial driver of gemcitabine resistance in PC.

Interestingly, whole-genome studies on PC patient samples have revealed mutations in the SLIT and ROBO genes (Aung et al., 2018; Biankin et al., 2012; Waddell et al., 2015). These studies have highlighted a strong correlation between the dysregulated SLIT/ROBO signaling and early stages of pancreatic neoplastic transformation (Biankin et al., 2012). The data from these studies strongly support the idea that aberrant regulation of axon guidance pathways is not merely a consequence but may actively drive the early events of tumor initiation and subsequent progression. Further, several independent studies have reported a crucial role of SLIT2, ROBO1 and ROBO2 proteins in PC initiation, progression and metastasis (Biankin et al., 2012; Chen et al., 2021; Göhrig et al., 2014; Han et al., 2015; Pinho et al., 2018). Moreover, ROBO3 was shown to drive PC metastasis and chemoresistance by utilizing patient-derived xenografts and FACS-sorted PC patient biopsies (Krebs et al., 2022). Notably, our unbiased transcriptomics data identified SLIT3, which was significantly decreased upon Orai3 silencing in GemR cells. Importantly, our earlier study has demonstrated that Orai3 is overexpressed in PC samples in comparison to healthy control pancreas and higher Orai3 expression is associated with poor prognosis (Arora et al., 2021). These observations provide a compelling mechanistic link where Orai3 may serve as a critical upstream modulator that initiates the activation of the SLIT/ROBO signaling pathway in PC progression. Thus, Orai3 may play a pivotal role in orchestrating the early molecular events that facilitate neoplastic transformation in pancreatic tissue, thereby contributing to PC progression, metastasis and chemoresistance.

NFAT transcription factor family has been widely studied in the context of immune cell function and disorders (Müller et al., 2010). Emerging literature reports elevated expression and enhanced transcriptional activity of NFAT isoforms in a range of cancers, which in turn augments the expression of genes driving tumorigenesis (Mancini et al., 2009). NFATc1-c4 are regulated by calcium-activated calcineurin and require sustained calcium signals, which are achieved through the opening of CRAC channels at the plasma membrane (Oh-hora et al., 2009). In our study, we employed an unbiased bioinformatic approach to explore the relation of NFAT transcription factor family with gemcitabine resistance in PC **(Fig. 8A,B)**. We observed that NFATc1 overexpression results in an increase in SLIT3 expression and promoter activity, suggesting that NFATc1 positively regulates SLIT3 transcription. Emerging evidence positions NFATc1 as the central regulator of tumor cell proliferation, survival, and metastatic progression (Mancini et al., 2009; Müller et al., 2010). An interesting study uncovered the contrasting roles for NFATc1 and NFATc2 in tumorigenesis, with NFATc2 functioning as a tumor suppressor while NFATc1 acting as an oncogene(Robbs et al., 2008). In PC cells, NFATc1 drives proliferation and anchorage-independent growth by activating c-myc transcription (Buchholz et al., 2006). Our findings reveal that NFATc1 transcriptionally regulates PC gemcitabine resistance by functioning as a bridge between Orai3 and SLIT3. Interestingly, our recent work demonstrated that NFATc1 plays an important role in regulating Orai3 expression in PC cells (Raju et al., 2024). Therefore, it emerges that NFATc1, Orai3 and SLIT3 work in intriguing feedforward loops to drive PC metastasis and chemoresistance.

Taken together, we reveal a crucial role of Orai3 in driving gemcitabine resistance. Further, we demonstrate that Orai3 regulates key characteristics of chemoresistance, such as enhanced proliferation, migration, invasion, stemness and metastasis. Moreover, through unbiased transcriptomics coupled with robust functional assays, we identify that SLIT3 functions downstream of Orai3 to drive gemcitabine resistance. Finally, we show that NFATc1 drives SLIT3 transcription thereby connecting Orai3 overexpression in GemR cells to SLIT3 augmentation **(Fig. 9)**. Collectively, this study suggests that Orai3 inhibition can enhance the efficacy of chemotherapy regimens. Therefore, future studies are required to identify specific drug molecules that inhibit Orai3. Eventually, these drugs may be administered in combination with gemcitabine to improve its therapeutic usefulness. This approach would offer a novel way to overcome chemoresistance in pancreatic cancer.

## Acknowledgements

This work was supported by the Anusandhan National Research Foundation (ANRF) project number SERB-CRG/2023/004054. RKM also acknowledges funding support from RCB Institutional Core funding, Department of Biotechnology, India (project number BT/PR52477/MED/30/2540/2024) and DBT/Wellcome Trust India Alliance Fellowship (IA/I/19/2/504651). The authors thank members of the Motiani laboratory for discussions and critical reading of the manuscript. SA acknowledges her Senior Research Fellowship from DBT, India. AT acknowledges his Junior Research Fellowship awarded from DBT, India. Illustrations were made using Biorender.

## Authors contribution

Samriddhi Arora: Methodology, Investigation, Visualization, Formal analysis, Writing- Original draft preparation. Gyan Ranjan: Investigation, Formal analysis, Visualization. Abhishek Tanwar: Investigation, Formal analysis, Visualization. Rajender K Motiani: Conceptualization, Supervision, Writing- Original draft preparation, Reviewing and Editing, Project administration, Funding acquisition.

## Supplementary Figures

**Supplementary Figure 1:**
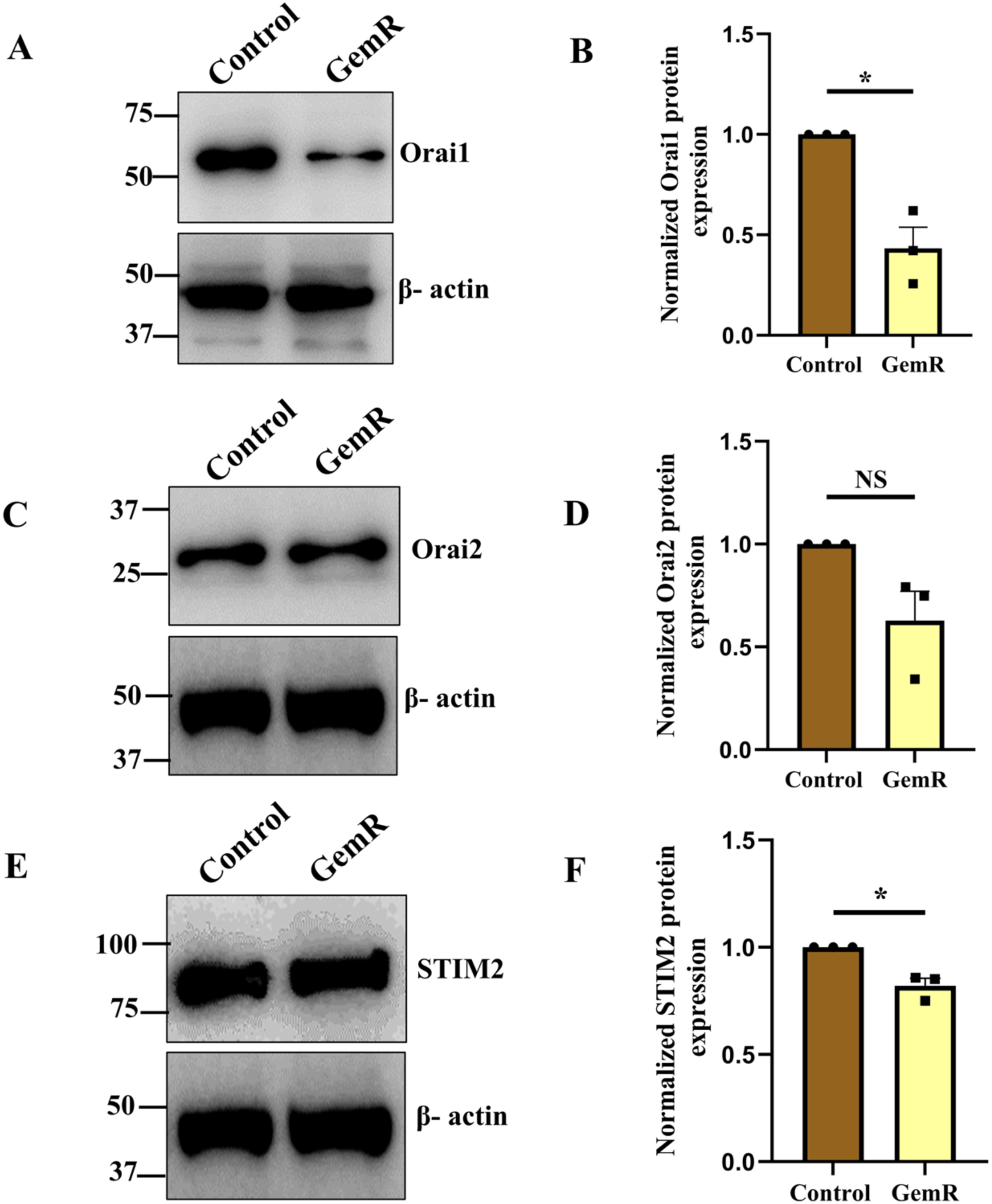
Expression of Orai1, Orai2 and STIM2 in CFPAC-1 GemR cells. **A)** Representative western blot showing Orai1 expression in Parental and GemR CFPAC-1 cells (N=3). **B)** Densitometric quantitation of Orai1 expression normalized to loading control β – actin. **C)** Representative western blot showing Orai2 expression in Parental and GemR CFPAC-1 cells (N=3). **D)** Densitometric quantitation of Orai2 expression normalized to loading control β – actin. **E)** Representative western blot showing STIM2 expression in Parental and GemR CFPAC-1 cells (N=3). **F)** Densitometric quantitation of STIM2 expression normalized to loading control β – actin. Data presented are mean ± S.E.M. A one-sample t-test was performed for panels B, D and F using GraphPad Prism software. Here, * *p* <0.05; ** *p* < 0.01; *** *p* < 0.001 and **** *p* < 0.0001.

**Supplementary Figure 2:**
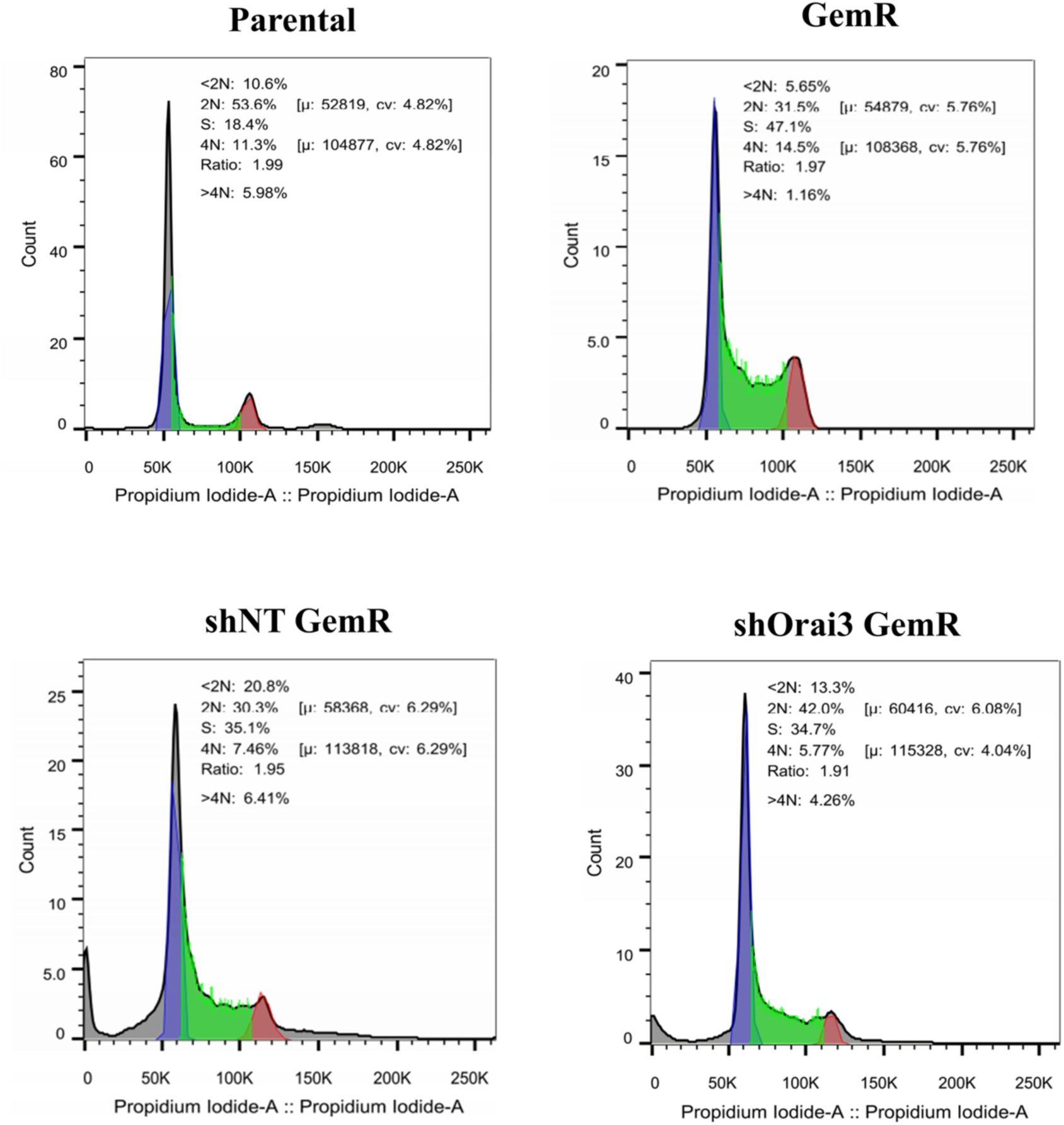
Orai3 regulates G1 to S transition of CFPAC-1 GemR cells. Representative cell cycle plots of parental, GemR, shNT GemR and shOrai3 GemR, depicting different cell cycle phases, generated using FlowJo software.

**Supplementary Figure 3:**
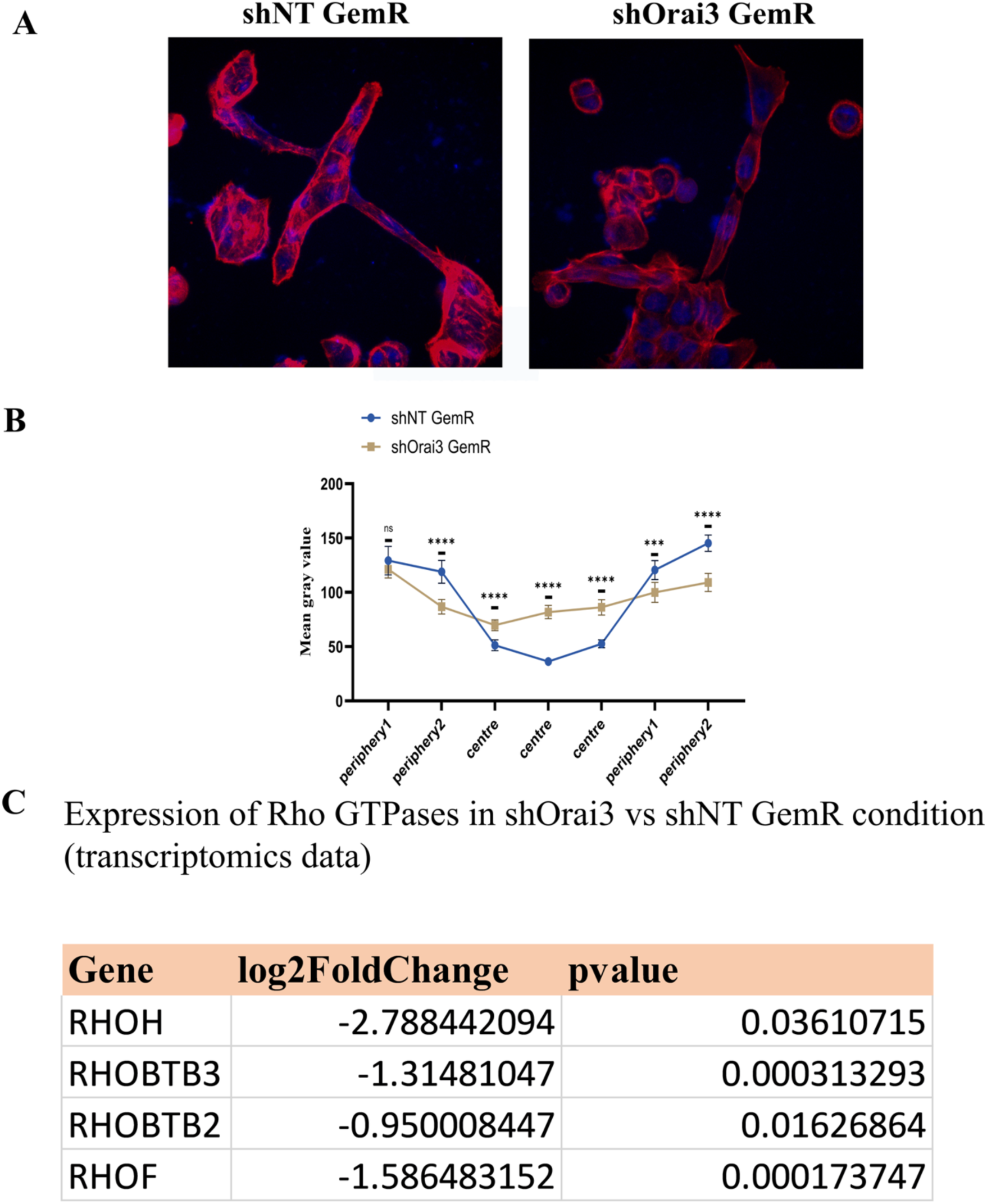
Orai3 regulates actin dynamics in gemcitabine-resistant cells. **A)** Representative confocal images show actin filament arrangement in Parental, GemR, shNT GemR and shOrai3 GemR CFPAC-1 cells. Phalloidin (red) was used to stain the actin cytoskeleton, and DAPI (blue) was used to stain the cell nucleus **B)** Quantitation of staining intensity, reflecting the F-actin distribution in Parental vs GemR and shNT vs shOrai3 GemR CFPAC-1 cells. **C)** RhoGTPases expression in shOrai3 GemR CFPAC-1 cells relative to the control, as analysed from RNA-seq data. Data presented are mean ± S.E.M. A multiple unpaired t-tests was performed for panel B using GraphPad Prism software. Here, * *p* <0.05; ** *p* < 0.01; *** *p* < 0.001 and **** *p* < 0.0001.

**Supplementary Figure 4:**
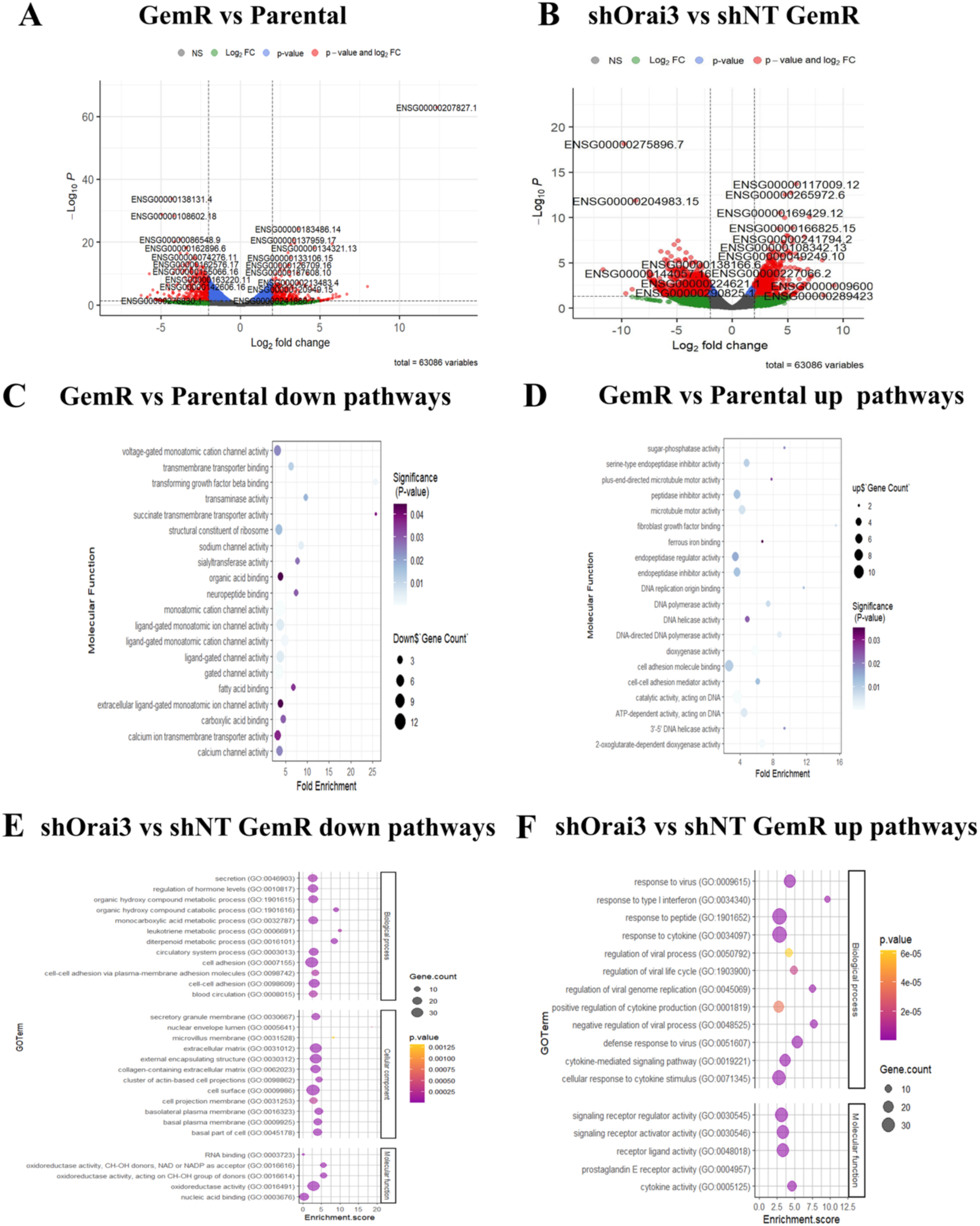
RNA-seq data analysis. **A)** Volcano plot generated after RNA-seq data analysis showing differentially regulated genes in GemR vs Parental cells. **B)** Volcano plot generated after RNA-seq data analysis showing differentially regulated genes in shOrai3 vs shNT CFPAC-1 GemR cells. **C)** GO pathway analysis of downregulated pathways in GemR vs Parental group. **D)** GO pathway analysis of upregulated pathways in GemR vs parental group. **E)** GO pathway analysis of downregulated pathways in shOrai3 vs shNT GemR. **F)** GO pathway analysis of upregulated pathways in shOrai3 vs shNT GemR.

## Materials and methods

### Cell culture

The cell line used in this study was obtained from the American Type Culture Center (ATCC). CFPAC-1 cells were grown in RPMI-1640 (Sigma) with 10% FBS, Panc-1 in DMEM high glucose (Sigma) with 10% FBS, and MiaPaCa-2 is also cultured in DMEM high glucose supplemented with 10% FBS and 2.5% horse serum (Himedia). Media used for culturing cells are also augmented with 1X anti-anti (Gibco) to prevent microbial contamination. Cells were maintained in a humidified chamber with 5% CO2 at 37^0^C.

### Western Blotting

The cells were washed with PBS (pH=7.4) and trypsinised using 0.1% trypsin to make cell pellets. The suspension was lysed using NP40 (Thermo Fisher) and a protease inhibitor cocktail overnight at -80^0^C. Protein lysates were collected by centrifugation at 13000 rpm at 4^0^C for 20min. Protein samples were quantified using a Bicinchoninic acid kit (Thermo Scientific). The quantified samples were loaded onto different concentrations of acrylamide gels for separation, followed by wet electrophoretic transfer to polyvinylidene difluoride (PVDF) membranes (0.45 µm; Millipore). The membranes were blocked with 5% skimmed milk powder (Himedia) in Tris-buffered saline with 0.1% Tween (TBST) for 1 hour at room temperature and probed with Orai3 antibody (1:500, Abcam), STIM1 antibody (1:500, BD biosciences), STIM2 (1:500, Abcam), Orai1 (1:500, Abcam), Orai2 (1:500, Abcam), E-cadherin (1:1000, Cell Signaling Technology), Vimentin (1:1000, Cell Signaling Technology), ALDH1A1 (1:1000, Cell Signaling Technology) and SLIT3 (1:100, SantaCruz) overnight at 4^0^C. The blots were then incubated with horse-radish peroxidase (HRP) conjugated secondary rabbit or mouse antibody for 2 hours at room temperature. The visualisation was performed using Image Quant LAS 4000 using a chemiluminescent substrate (Millipore).

|  | Brand | Catalog number |
| --- | --- | --- |
| Orai1 | Abcam | ab86748 |
| Orai2 | Abcam | ab180146 |
| Orai3 | Abcam | ab254260 |
| STIM1 | BD biosciences | 610954 |
| STIM2 | Abcam | ab157667 |
| ALDH1A1 | Cell Signalling Technology | 54135S |
| SLIT3 | SantaCruz | sc-293463 |

### RNA extraction

According to the manufacturer’s protocol, total RNA was extracted from cells using an RNAeasy mini prep kit (Qiagen). RNA was dissolved in 30 µl nuclease-free water. RNA extracted was quantified using a Nanodrop spectrophotometer (Thermo Fisher Scientific).

### RNA sequencing

Total RNA of different conditions parental, GemR, shNT and shOrai3 GemR were sent to Clevergene, Bengaluru (India) for RNA-sequencing. The sequenced data was generated using Illumina NovaSeq. The data quality was assessed using FastQC and MultiQC software. The quality control approved reads were aligned to the indexed human genome (GRCh38.p14) utilising the STAR v2 aligner. Gene-level expression values were derived as read counts employing feature-counts software. The DEseq2 package was used to analyze differential expression, and normalized counts were obtained. Data quality was assessed using FastQC and MultiQC software. GRCm39 was used as the reference genome. Normalized gene expression values were obtained from raw read counts using feature counts software. The differentially expressed genes were screened using log2FC of +2 and -2 for upregulated and downregulated genes, respectively. These genes were then analyzed for pathway enrichment analysis using GO pathway analysis software, and a bubble plot was generated to represent the enriched pathways. Common genes that were differentially regulated in GemR vs Parental and shOrai3 vs shNT were determined using a Venny 2.0 tool.

### Real-Time PCR

Extracted RNA was reverse transcribed to generate first-strand cDNA (High-Capacity cDNA Reverse Transcription Kit, ThermoFisher Scientific). Quantitative real-time PCR was performed using TB Green Premix Ex Taq II (SYBR green, Takara) according to the manufacturer’s instructions on a QuantStudio 6 flex real-time PCR system (ThermoFisher Scientific).

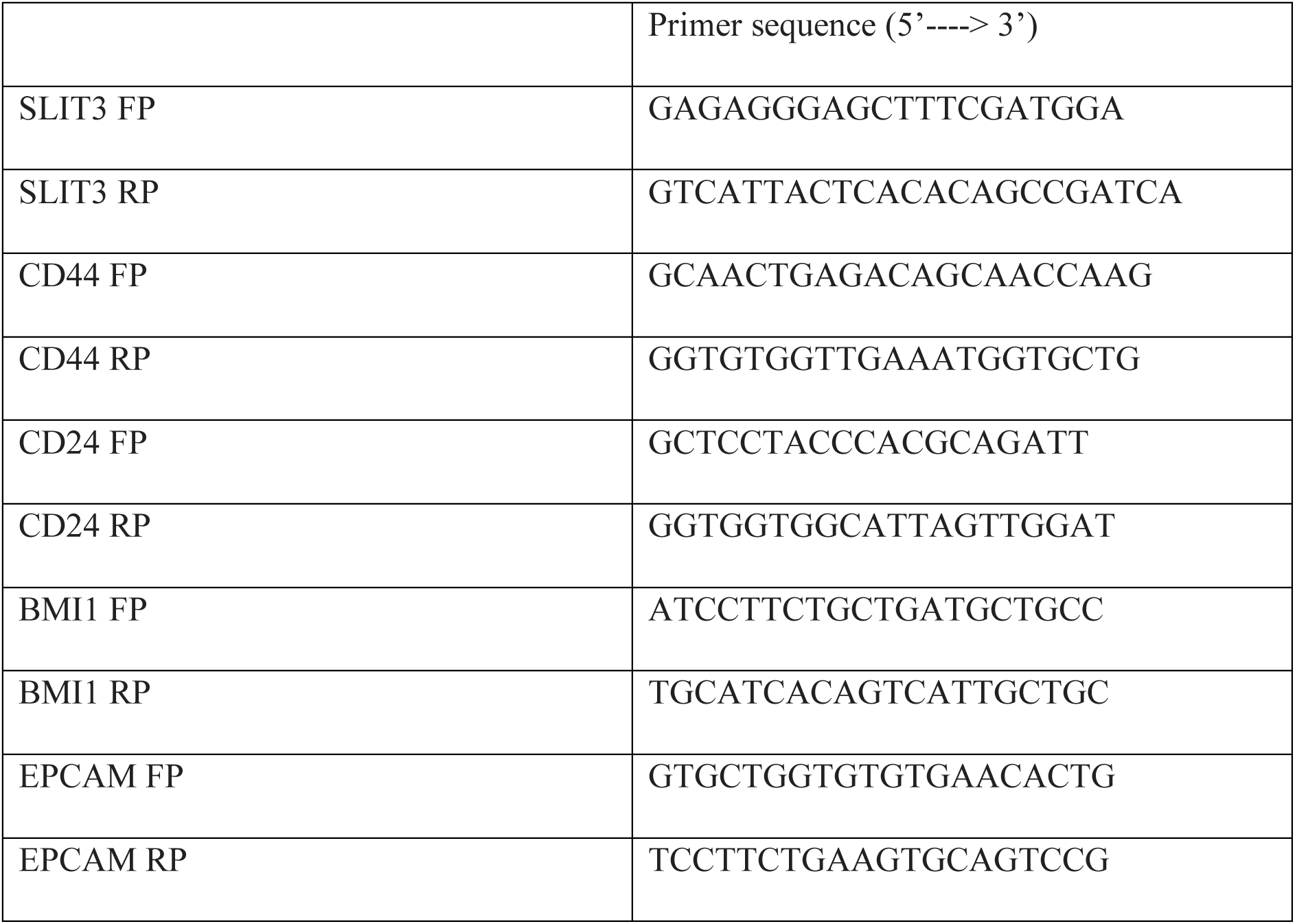

### SLIT3 promoter cloning

The human Slit3 promoter was obtained from Eukaryotic Promoter Database (EPD) and the sequence was analysed using NCBI-BLAST to find mRNA start site and the first codon. Primers were designed to amplify the 991 bp region (-969 to +23 w.r.t start codon) of the Slit3 core promoter. PCR amplification of the Slit3 promoter was done using Phusion High Fidelity Polymerase (F503, Thermo Fisher Scientific) and the PCR amplicon was cloned into psiCHECK-2 dual-luciferase reporter vector (C8021, Promega) at the SacII/MluI sites. The clones were verified through restriction digestion. The primers used for cloning are listed below:

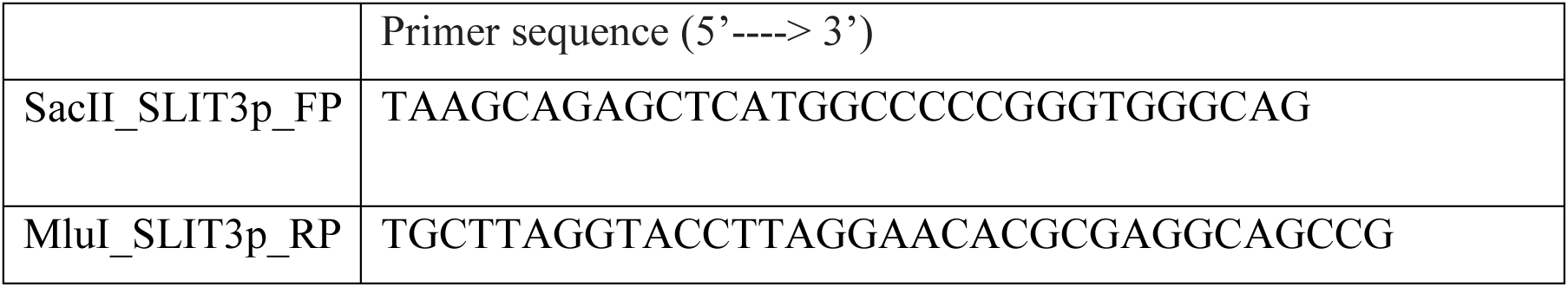

### Luciferase assay

CFPAC-1 GemR cells were seeded in a 24-well plate at a density of 50,000 cells/well. Cells were co-transfected with SLIT3p and siNT/siNFAT2 using a JetPrime transfection reagent (101000046, Sartorius). After 48h of transfection, cells were assayed for luciferase activity using the dual luciferase assay kit (E1910, Promega) as per manufacturer’s protocol. Renilla luciferase was used for normalization.

### Cell cycle analysis

To assess cell cycle progression, pancreatic cancer cells were seeded in a 6-well plate and treated with 5µM gemcitabine the next day. After 72 hr of treatment, cells were harvested and kept in 70% ethanol for 24 hours at -20^0^C. The next day, cells were treated with RNase at 37^0^C for 1 hour, then stained with Propidium Iodide (PI) in the dark for 30 minutes. The cell cycle distribution was detected using BD FACSVerse Cell Analyzer (BD Bioscience).

### Ca^2+^ Imaging

Pancreatic cancer cells were grown in confocal dishes (200350, SPL life sciences) for Ca^2+^ imaging experiments. Ratiometric dye Fura-2 AM (F1221, Invitrogen) was a Ca^2+^ indicator. Cells were loaded with Fura-2 AM for 30 minutes at 37^0^C and 5% CO2 in the corresponding media. The detailed experimental protocol was followed, as reported previously (Arora et al., 2021).

### Apoptotic assay

CFPAC-1 cells were seeded in a 24-well plate and treated with different concentrations of gemcitabine for 72 h. Cells were stained with Annexin V – TRITC labelled (AAT Bioquest) and DAPI to measure the cell death using BD FACSVerse Cell Analyzer (BD Bioscience).

### Spheroid formation assay

Spheroids were generated using the hanging drop method. A drop of 10 microlitres of complete media with 40,000 cells was pipetted onto the lid of 100mm Petri dishes and was inverted over the dish containing PBS to avoid drying. Dishes were stored at 37^0^C for four days for spheroids to form. After the period of incubation, images were taken using a Nikon stereo zoom microscope, and the area of spheroids was calculated using NIS Elements D version 5.21.03 software.

### Migration assay

To study migration assay, we used Boyden chamber transwells (3422, Corning). Different groups (Parental, GemR, shNT, and shOrai3 GemR) of CFPAC-1 cells (1 × 10^5) were seeded in the medium containing 3% FBS in the upper chambers of the 24-well plate. The lower chambers were filled with 650 ul of medium containing 10% FBS. The cells were incubated for 48 h at 37^0^C. After the incubation period, cells were fixed with 3.7% formaldehyde, permeabilized using 100% methanol, and stained with 1% solution of crystal violet. The migrated cells were counted using ImageJ software.

### Cell proliferation assay

Cell proliferation was assessed using a BrdU-based colorimetric immunoassay kit (11647229001, Roche). 5 × 10^3 cells were seeded for different groups (Parental, GemR, shNT, and shOrai3 GemR) in 96-well plates, and the proliferation rate was measured at various time points. The steps in quantifying cell proliferation were followed as per the protocol provided with the kit.

### Phalloidin staining

For actin filament visualization, cells were grown on a glass coverslip. After 24 hours, cells were washed thrice with chilled PBS, then fixed on ice with 3.75% formaldehyde solution in PBS for 15 min. Following fixation, cells were permeabilized with 200 µl 0.5% Triton X-100 at room temperature for 10 min. Blocking was done using 1% BSA in PBS at 4^0^C for 30 min. Cells were stained with Alexa Fluor 594 Phalloidin (Thermofisher Scientific, A12381) for 20 min at room temperature. After staining, coverslips were air dried, followed by mounting with SlowFade™ Gold Antifade Mountant with DAPI (ThermoFisher Scientific, S36938) and imaged using a confocal microscope. F-actin distribution was quantified using ImageJ software, according to the published methodology (Zonderland et al., 2019).

### Zebrafish husbandry

The zebrafish used in this study were housed at the Regional Center for Biotechnology, India, in compliance with standard ethical protocols approved by the Institutional Animal Ethics Committee. All efforts were made to minimize animal suffering.

### Zebrafish xenograft microinjection

Microinjections were performed using the human pancreatic cancer cell line CFPAC- 1(Parental, GemR, shNT GemR and shOrai3 GemR). Cells were stained with lipophilic Vybrant CM-DiI (ThermoFisher Scientific) at a 4 µL/mL concentration. The staining procedure was performed as per the manufacturer’s instructions. Cells were resuspended to a final concentration of 30x 10^3^/µL in 1X Versene (0.2 g EDTA(Na4) per liter of Phosphate Buffered Saline) before injection to avoid capillary clogging. The labelled tumor cells were microinjected into zebrafish embryos using borosilicate glass capillaries under a fluorescence stereomicroscope (Nikon, SMZ800N) equipped with a FemtoJet 4i microinjector (Eppendorf). Cells were injected into the perivitelline space (PVS) of 2-day-post-fertilization (pdf) zebrafish embryos, which were pre-anesthetized with 1X Tricaine (Sigma-Aldrich). Following injection, the zebrafish xenografts were transferred to E3 medium and incubated at 34 °C. At 1-hour post-injection (hpi), the xenografts were examined for the presence or absence of cells in circulation, and those with severe oedema, cells in the yolk sac, cell debris, or noninjected were excluded (Martinez-Lopez et al., 2021). At four dpi, xenografts were anaesthetized to image the metastatic cells in caudal hematopoietic tissue (CHT). Xenografts were kept in 1X Phenylthiourea (PTU) to suppress pigmentation until the experiment was completed.

### Statistical Analysis

All the experiments were performed as three or more independent biologicals. Data shown as mean ± SEM. A one sample t-test and unpaired student’s t-test were performed to determine the statistical significance. A one-way ANOVA was performed for data containing more than two conditions. p < 0.05 was considered as significant and is presented as ‘‘*’’, p <0.01 is presented as ‘‘**’’, p < 0.001 is presented as “***” and p <0.0001 is presented as “****”.

## Data Availability

The raw sequencing data will be made available upon request. Additional supporting data, not included in the main figures, is provided in the supplementary file.

## Conflict of Interest Disclosure

The authors declare no conflict of interest.

## Notes

### Competing Interest Statement

The authors have declared no competing interest.

